# Dissecting the binding mechanisms of transcription factors to DNA using a statistical thermodynamics framework

**DOI:** 10.1101/666446

**Authors:** Patrick C.N. Martin, Nicolae Radu Zabet

## Abstract

Transcription Factors (TFs) bind to DNA and control activity of target genes. Here, we present ChIPanalyser, a user-friendly, versatile and powerful R/Bioconductor package predicting and modelling the binding of TFs to DNA. ChIPanalyser performs similarly to state-of-the-art tools, but is an *explainable model* and provides biological insights into binding mechanisms of TFs. We focused on investigating the binding mechanisms of three TFs that are known architectural proteins CTCF, BEAF-32 and su(Hw) in three Drosophila cell lines (BG3, Kc167 and S2). While CTCF preferentially binds only to a subset of high affinity sites located mainly in open chromatin, BEAF-32 binds to most of its high affinity binding sites available in open chromatin. In contrast, su(Hw) binds to both open chromatin and also partially closed chromatin. Most importantly, differences in TF binding profiles between cell lines for these TFs are mainly driven by differences in DNA accessibility and not by differences in TF concentrations between cell lines. Finally, we investigated binding of Hox TFs in Drosophila and found that Ubx binds only in open chromatin, while Abd-B and Dfd are capable to bind in both open and partially closed chromatin. Overall, our results show that TFs display different binding mechanisms and that our model is able to recapitulate this diverse repertoire of mechanisms.

## Background

Decades of research have shown that gene expression is at the heart of many, if not all, cellular processes. From development to cellular homoeostasis, the activation or repression of gene expression enables cells, and by extension organisms, to function properly. One of the key components of the regulation of gene expression is Transcription Factors (TFs). TFs are a class of proteins that bind to DNA in a sequence specific manner [1, 2]. The most commonly used experimental method to determine specific regions of DNA where TFs bind is chromatin immunoprecipitation followed by sequencing (ChIP-seq) [3, 4]. This technique has become the gold standard to determine the binding profiles of TFs to the genome, but, despite the huge impact on understanding gene regulation, it does not provide a mechanistic model of what drives the binding of TFs to those regions or even how genes are regulated. While we still lack a complete predictive model for gene expression, over the years, many factors have been identified as contributing to context dependant TF binding.

The most fundamental aspect to consider concerning TF binding specificity is the DNA sequence itself. Most TFs have a preferred binding motif [5, 6]. The most common way to describe this motif is in the form of a Position Weight Matrix (PWM); a measure of binding energy between TFs and DNA weighted by the genomic base pair frequency [5, 7]. Nevertheless, TFs can have tens of thousands of potential binding sites within each genome, yet only a few hundred will be occupied by TFs [8, 9].

Previous studies have shown that some TF binding events are TF concentration dependent [10, 11, 12, 13], where varying the concentration of the TF will drive the expression of different sets of genes. However, there are many more spurious sites, rather than functional binding sites where TF’s could bind. This still begs the question: how do TFs distinguish between bound and unbound binding sites?

One way to reduce the number of available sites is to consider DNA accessibility. Are these sites even available for binding in the first place? This assumes that TFs would bind only to sites that are accessible and cannot locate sites within dense chromatin [14, 15]. Nevertheless, there is a certain class of TFs known as pioneer TFs are able to bind in closed chromatin. More specifically, pioneer TFs can bind sites in closed dense chromatin and subsequently open the chromatin [16, 17, 18, 19].

Over the years, many tools and frameworks have aimed to predict transcription factor binding. One of the earliest tools incorporating DNA accessibility was the PIQ algorithm (Protein Interaction Quantification) which implemented a machine learning type algorithm to filter out binding sites located in inaccessible DNA [20]. Later, msCENTIPEDE improved upon CENTIPEDE using multi-scale models for inhomogeneous Poisson processes to untangle TF binding with respect to DNA accessibility [21]. Some notable tools that have been developed through DREAM challenges are FactorNet, implementing a deep learning framework [22], Anchor, relying on a XGBoost system [23] and Catchitt making use of supervised machine learning and iterative training [24]. While machine learning methods predict TF binding events with high accuracy, they are often difficult to interpret. These methods are not always the best suited to understand the mechanism driving a biological phenomenon.

We previously showed that statistical thermodynamics can be used to model TF binding to DNA with high accuracy [13]. Considering only binding energy between TFs and DNA (estimated by the PWM and a scaling factor modulating the binding energy), the number of bound molecules to the DNA and DNA accessibility, we modelled binding of five TFs in Drosophila embryo. Our results confirmed that, for some TFs, this model is sufficient to explain the majority of observed binding events in ChIP data and we were able to backwards infer number of bound molecules and specificity for five TFs in Drosophila embryo (bcd, cad, gt, hb and Kr).

In this manuscript, we build upon our previous model and developed ChIPanalyser a versatile and user-friendly R/Bioconductor package [25, 26]. Furthermore, we used this model to describe the behaviour of several Drosophila TFs (CTCF, BEAF-32, su(Hw), Ubx, Abd-B and Dfd) in different cell lines (BG3, Kc167 and S2).

## Methods

### Model Description

ChIPanalyser is an R package available on Bionconductor [26]. The package is an implementation of the statistical thermodynamics model proposed in [13]. Briefly, the model requires *(i)* a PWM (Position Weight Matrix) or PFM (Position Frequency Matrices) of the TF of interest, *(ii)* DNA accessibility data to model binding site accessibility and two additional parameters: *(iii) λ* (a PWM scaling factor) and *(iv) N* (the number of bound molecules) [13]. The probability of a position *j* on the DNA being occupied is given by [13]:

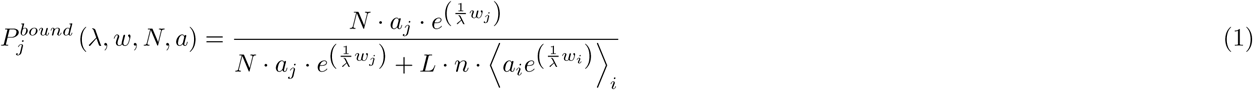

*λ* and *N* are difficult to estimate from experimental data and, thus, we used ChIP-seq data and selected the values of these parameters that maximise (or minimise) the goodness of fit metrics. *L* represents the length of the genome and *n* the ploidy level of the organism.

### Datasets

To carry out the analysis described in this manuscript, we selected data originating from various sources (see Table S1)

#### DNA Sequence

Reference Sequences of *Drosophila melanogaster* (dm6) and *Homo sapiens* (hg38) were extracted from the Bsgenome R packages. All data sets were either aligned to the dm6 versions of the Drosophila genome or lifted over from dm3 to dm6 using the UCSC genome liftover chain.

#### PWM and PFM

Binding Motif matrices were downloaded from online repositories such as JASPAR [27] or extracted from the MotifDb R package [28], which collects and compiles PFMs and Position Probability Matrices (PPM) from various online repositories (see Figure S1 in *Supplementary Materials*). For the purpose of method comparison requirements (msCENTIPEDE), TF binding sites were extracted using FIMO from the MEME-suit tool kit [29].

#### ChIP-seq

ChIP enrichment signal and ChIP peaks were downloaded (pre-processed) from modEncode in three Drosophila cell line: Kc167, S2 and BG3. For the purpose of this study, we considered ChIP-chip and ChIP-seq as sufficiently similar to be comparable as ChIPanalayser focuses on ChIP signal pile up for parameter inference and goodness of fit. When it was required, supplementary data sets were downloaded from GEO. GEO datasets were aligned to the genome (dm6) using bowtie-2 (- -non-deterministic). *SAM* files were converted to *BAM* files using samtools [30]. Peaks and pile-up signal were called using macs2 with a 0.01 FDR (-q 0.01) [31]. Processed data sets for *Homo sapiens* were directly downloaded from ENCODE were already aligned to hg38. We selected one of the data sets provided and used in the DREAM challenge competition related to TF binding prediction. When required, peak replicates were combined using the GenomicsRanges package in R. Datasets used for this analysis are described in Table S1 in *Supplementary Materials*.

#### DNA accessibility

DNase I hypersensitivity data was generated by modEncode for the three cell lines used in this analysis [32]. We aligned fastq files to the dm6 genome build using bowtie-2 (- – non-deterministic). *SAM* files were converted to *BAM* files using samtools [30]. Peaks and read pile-ups were called and produced using macs2 (–broad-call -cutoff 0.05 -q 0.05) [31]. DNase I hypersensitvity data for *Homo sapiens* was directly downloaded from ENCODE and replicates were merged using samtools. When required, DNase peak replicates were combined using the GenomicsRanges package in R. The level of accessibility is consistent with past experiments (see Figure S2). ATAC-seq data for Kc167 cells was used from [33] and ATAC-seq scores were computed using macs2 as described in [33]. We selected a series of ATAC-seq signal thresholds that we would use as a cut-off point to select accessible/inaccessible DNA. These thresholds were based on signal quantiles from 0.05 to 0.95 by 0.05. We also considered 0.99, 0.999, 0.9999 quantile thresholds. We will refer to this method as Quantized Density Accessibility (QDA).

#### RNA-seq

In order to rescale TF abundance between cell lines, we used RNA-seq data from [34]. RNA-seq relative abundance was used to rescale the estimated number of bound molecules from one cell line to another.

### Description of ChIPanalyser

The workflow of ChIPanalyser is described in Figure 1. Briefly, the optimal set of parameters (for *λ* and *N*) can be inferred from ChIP data by maximising (or minimising) a goodness of fit metric. Nevertheless, if the user approximates these values by other means, ChIPanalyser does not require any training data at all. Using these values, ChIPanalyser will produce base pair resolution ChIP like profiles for different genomic regions and compare the prediction with the actual ChIP data (if that is provided by the user).

**Figure 1:**
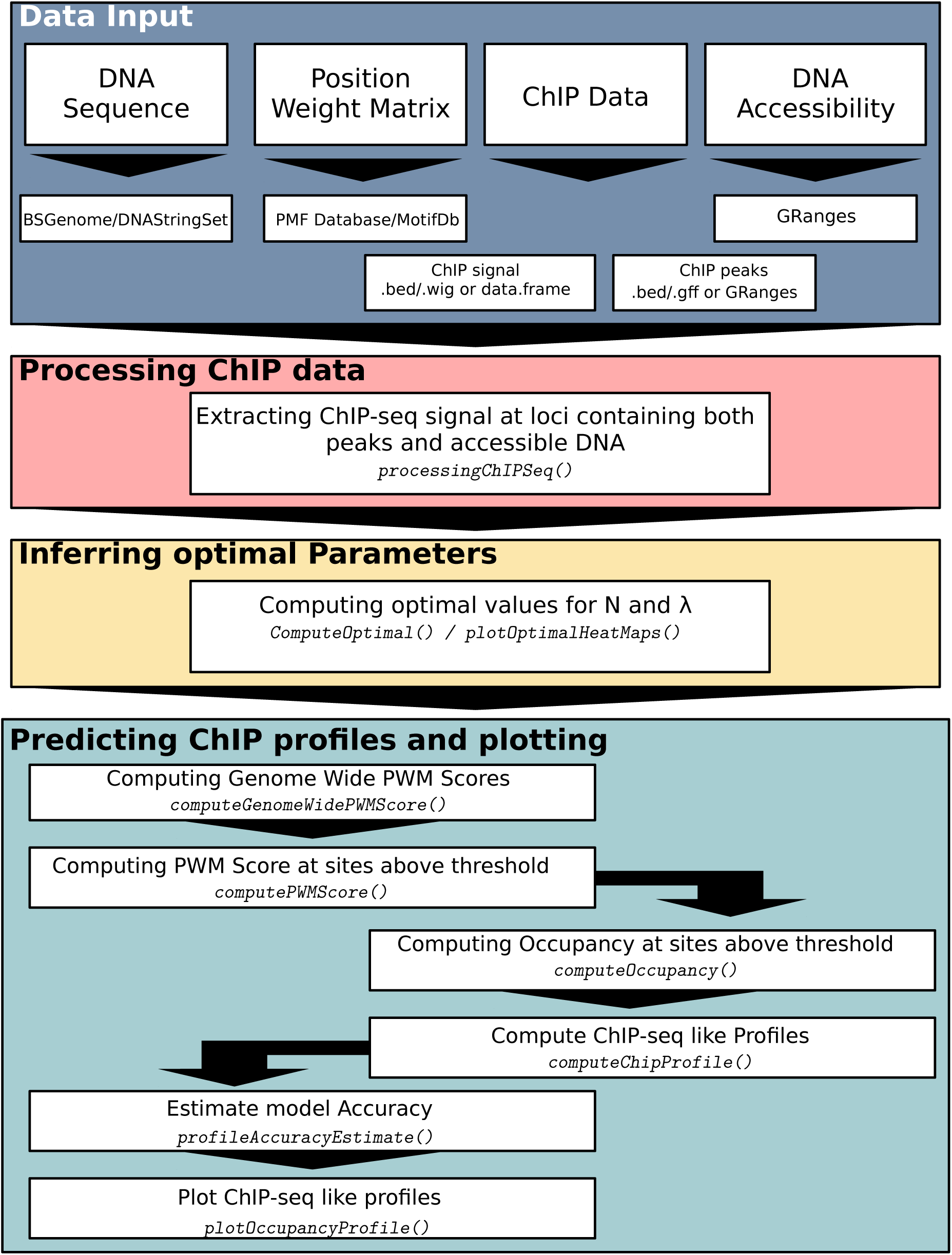
ChIPanalyser workflow. ChIPanalyser follows the following work flow. **Data Input:** Data may come in various formats (e.g. bed, wig, gff etc.). **Processing ChIP-seq data:** If ChIP data is used to infer the optimal set of parameters (and/or validate model goodness of fit), ChIP data will be normalised and only regions of interest will be extracted for further analysis. **Inferring optimal parameters:** Inferring optimal parameters will be achieved by maximising (or minimising) a goodness of fit metric. **Predicting ChIP profiles and plotting:** Using the optimal values for number of bound molecules and the PWM scaling factor, ChIPanalyser will produce ChIP like profiles. Both optimal parameter heatmaps and ChIP profiles can be plotted using the package’s plotting functions.

ChIPanalyser uses a set of genomic regions to infer optimal parameters. If the genomic regions are not provided by the user, the top *n* regions will be selected based on highest ChIP score after binning the genome into bins of 20 Kb (number of regions to be selected and bin width and can be customized). For our analysis, we split the entire genome into bins of 20 Kb and selected bins that contained at least one CTCF, BEAF-32 and su(Hw) peaks in at least one biological replicate and at least one cell line. By doing so, we ensure that the regions we will use in this analysis are common between all data sets. This resulted in 3293 bins of 20 Kb that contain at least one peak of any of these architectural proteins and at least one base pair of accessible DNA. In addition, we followed the same process for Hox transcription factors, which resulted in a total of 3838 bins of 20 Kb containing at least one peak for each TF (Ubx, Abd-b, and Dfd). Normalised and ordered bins (based on highest ChIP scores in that bin) were produced by the *processingChIP* function provided by ChIPanalyser. Following this, we selected the top ten regions in order to train our model (to infer N and *λ* by maximising or minimising a goodness of fit metric). The top ten regions contain the strongest peaks which ensures that we have a balance between True Positive and True Negative signals. Once we had selected the optimal set of parameters based on our training set, we validated our results on the other regions that do not contain the training set.

During this step of the analysis (*processingChIP*), we also included a noise filtering method. The current model does not consider ChIP depletion, therefore all negative scores are replaced by 0. With that in mind, ChIPanlyser provides four methods of filtering noise: *Zero, Mean, Median* and *Sigmoid. Zero* removes only depletion scores (equivalent to “no noise filtering”). *Mean* and *Median* replace all scores below the mean and the median after filtering out depletion scores. Finally, *Sigmoid* applies a logistic weighting to every score, modulating ChIP scores around the 95th quantile point. All analysis in this manuscript was carried out after using the *Sigmoid* noise filtering method.

Once the *loci* of interest have been selected, we inferred the optimal set of parameters by using ChIPanalyser’s *computeOptimal* function. The optimal set of parameters are inferred by maximising (or minimising) the average goodness of fit metric over all regions selected. ChIPanalyser offers 12 different metrics: correlation coefficients (Pearson, Spearman and Kendall), Mean Squared Error (MSE), Kolmogorov-Smirnov Distance, precision, recall, accuracy, F-score, Matthew’s correlation coefficient (MCC) and Area Under Curve Receiver Operator Characteristic (AUC ROC or just AUC) (see Table S2 in Supplementary Materials). We also developed a novel method that describes the ratio of shared geometric area between curves and difference in area between curves. ChIPanalyser generates a ChIP like profile at a base pair level resolution, however window size may be adjusted. The goal is to mimic experimental ChIP profiles by smoothing high occupancy binding sites into ChIP like profiles. This approach was described by [13].

For this analysis we used a 100 bp window for validation. Goodness of fit is carried out by comparing our prediction to ChIP score data (as opposed to peak location overlap). The rationale behind using ChIP scores instead of peaks was two-fold: *(i)* we consider peaks that are missed by peak calling algorithms and *(ii)* using ChIP scores ensures that we also consider signal enrichment. The latter is particularly relevant when estimating the number of bound molecules.

The evaluation method used by ChIPanalyer is significantly more stringent than methods used in other frameworks. When confusion matrices are required for scoring (AUC, recall, F-score, MCC, Accuracy, precision), ChIPanalyser uses 20 threshold values bound between the lowest occupancy score (predicted or experimental score) and the highest occupancy score (predicted or experimental score). The threshold values are squared in order to ensure a higher density of threshold values close to the lower end of occupancy scores. For every threshold value, ChIPanalyser compares its predicted profile to the experimental profile in 100 bp bins. If they both contain a “signal”, we consider that ChIPanalyser has correctly predicted local ChIP enrichment. If ChIPanalyser predicts ChIP enrichment when no experimental signal is present, we consider this bin to be a false positive case. The same approach was used for false negative cases (Experimental enrichment but no predicted enrichment) and true negative cases (No enrichment in either experimental or predicted profiles). This approach ensures that the model is penalised if it fails to predict peak enrichment or conversely over estimates peak enrichment.

The optimal parameters inferred over training can be visualised in the form of a heatmap describing the score associated to each combination of *λ* and *N*. Heatmaps are produced using the *plotOptimalHeatMaps* function. Finally, using the optimal set of parameters, ChIPanalyser will produce ChIP like profiles that can be visualised using the *plotOccupancyProfiles* function provided by the package.

## Results

### Evaluation of ChIPanalyser

Previously, we showed how statistical thermodynamics can be used to mechanistically explain the binding of TFs in *Drosophila* [13]. The optimal set of parameters (see Methods) was inferred by maximising correlation and minimising Mean Squared Error (MSE) between the predicted profile and experimental ChIP data. Nevertheless, we observed that, in some cases, the predicted profiles and ChIP profiles display low correlation coefficient despite the profiles looking similar and vice versa (e.g. see Figure S3A and S3B in *Supplementary Figures*). In some cases, selecting the optimal parameters was hindered by little variation in correlation between parameter combinations and, thus, the selection of these parameters was exclusively driven by MSE (see Figure S3C in *Supplementary Figures*).

To reduce the potential influence of background noise, we tested four noise removal methods: *Zero, Mean, Median* and *Sigmoid*; see Methods. To test the performance of these methods, we used three CTCF datasets (see Table S1 in *Supplementary Tables*): *(i)* a ChIP-chip dataset with very little background noise (modEncode 2639), *(ii)* a ChIP-seq dataset with high background noise (modEncode 3674) and *(iii)* a combination of all ChIP datasets in S2 cells (by adding enrichment signals together at a base pair level). To ensure equal contribution of each data set, we normalised the signal. We ran the model on the top ten regions (as described in Methods) and searched for the optimal set of parameters (*λ* and *N*) that optimised the goodness of fit metric (in this instance – AUC). All four noise filtering methods have little to no effect on ChIP data (see Figure S4 in *Supplementary Figures*). The Sigmoid method showed a slight signal reduction in smaller peaks (especially for noisy datasets), which was then translated into a slight improvement of the mean Area Under Curve Receiver Operator Characteristic (AUC ROC) score between ChIP signal and our predictions (see Figure S4 in *Supplementary Figures*).

In addition to Pearson correlation and MSE, we tested several goodness of fit metrics to verify the influence of the metrics on our model as described in Methods and Table S2 in *Supplementary Tables*. We used the same three CTCF datasets as described above and observed the emergence of two classes within these metrics: *(i)* similarity metrics that describe how similar the two curves are (correlation coefficients, precision, MCC, Accuracy, F-score and AUC ROC) and *(ii)* dissimilarity metrics that are a measure of how different two curves are (MSE, geometric ratio, recall and Kolmogorov-Smirnov distance). Our results showed that depending on the metric used, the optimal set of parameters varied significantly, but each of the two classes (similarity and dissimilarity metrics) displayed similar yet not identical values for the optimal parameters (see Supplementary Figure S5 A-F).

Goodness of fit metrics influence the way the model selects the optimal parameters, but how does this translate to the individual predicted ChIP profile level? We further investigated this behaviour at the individual *loci* using the same three CTCF datasets. Figure 2A-B shows that similarity metrics (black shades) tend to be less prone to false positive peaks but miss the actual ChIP signal strength within the peak (the height of the peak). On the other hand, dissimilarity metrics (light blue shade) generate far more false positives but accurately recover the height of the peaks.

**Figure 2:**
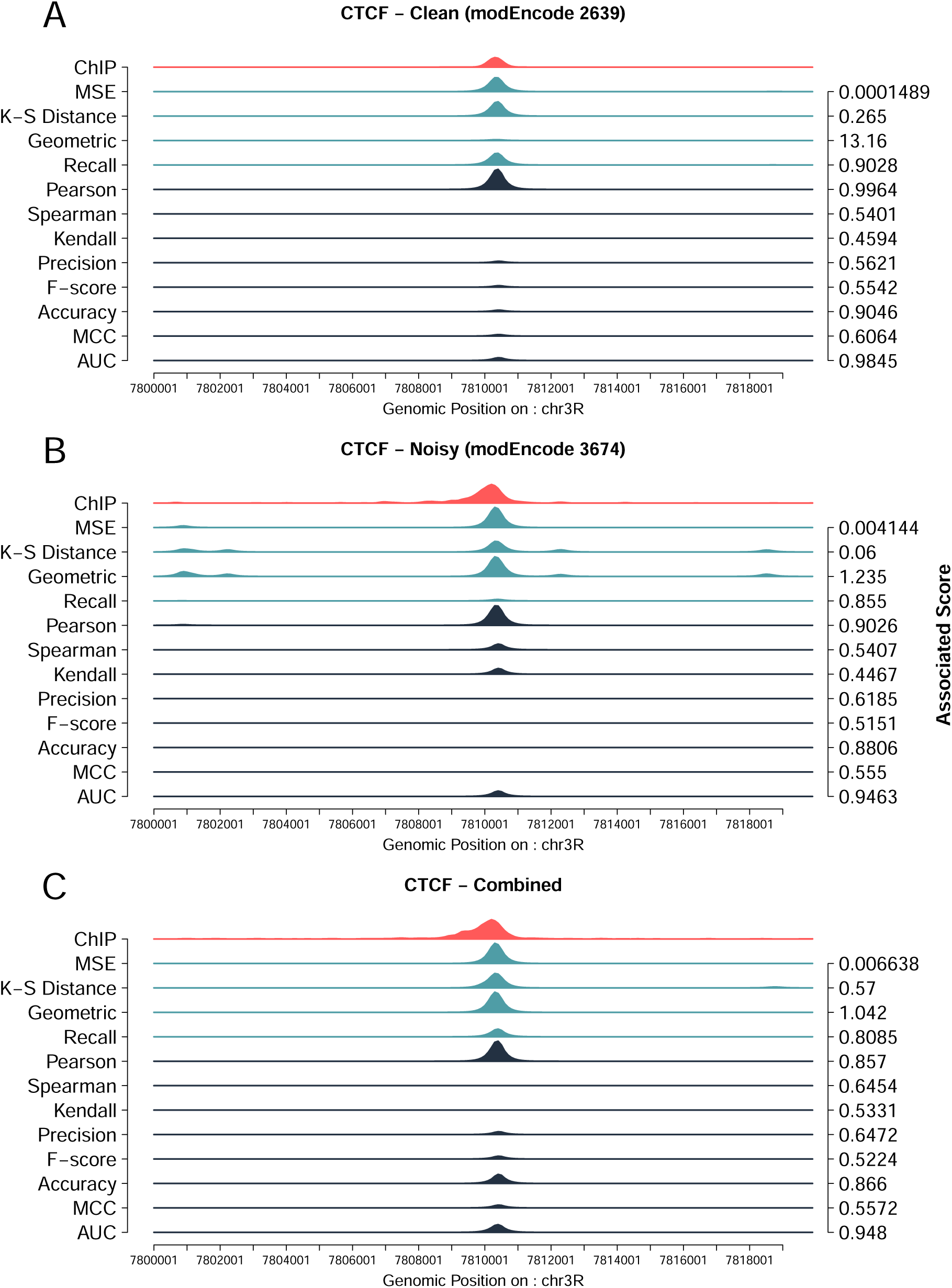
Goodness of fit Methods are context dependent. (**A**) ChIPanalyser correctly predicts CTCF peaks in a clean ChIP dataset (modEncode 2639) for the majority of metrics used. (**B**) For a noisier dataset (modEncode 3674), dissimilarity metrics capture the height of the peak but also tend to show a high rate of False Positive peaks. In contrast, similarity metrics accurately predict the location of the peak, but tend to underestimate peak height. (**C**) Combining several ChIP replicates (all ChIP-seq datasets in S2 cells; see Table S1 in Supplementary Materials) does not reduce the rate of False Positive peaks for similarity metrics. The red profile shows experimental ChIP peaks, while light blue and dark blue are predicted profiles. Light blue and dark blue as dissimilarity and similarity metrics respectively. Associated scores are the scores for each profile when that metric was used to select optimal parameters.

Overall, the best performing metrics were AUC ROC and MSE. AUC ROC occasionally missed peak enrichment completely however, seemed to recover peak location fairly accurately, while MSE rarely missed peak enrichment but also produced a higher number of false positive peaks. For much of the following analysis, we used AUC ROC and MSE, since they are more widely used estimators and performed best. More specifically, MSE was used as the training metric to select the optimal set of parameters. AUC, recall, Spearman correlation and MSE were used for validating model performance.

To evaluate the performance of our model, we first used a chromosome withholding set up. The model was trained on the top 10 regions (as described in Methods) on chromosome 3R (Figure 3A). We then validated our model using two approaches: *(i)* on the top 10 regions found on chr2R (Figure 3B) and *(ii)* on the top 10 regions on chr3R excluding regions used for training (Figure 3C). Our results show that ChIPanalyser accurately recovers peak location and peak enrichment between chromosomes (Figure 3D-G).

**Figure 3:**
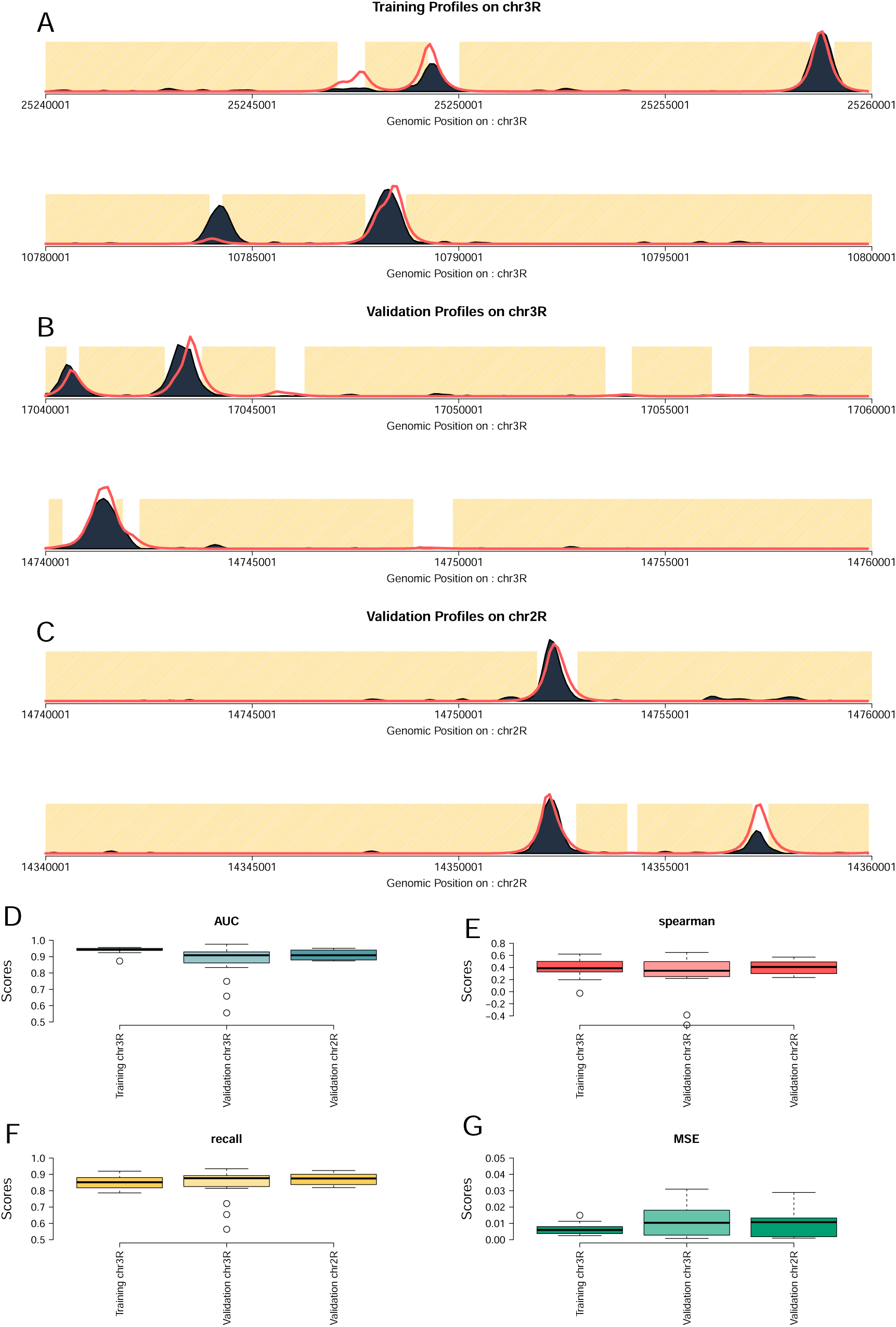
Chromosome withholding setup for model validation. We analysed BEAF-32 ChIP in S2 cells (modEn-code 922) and we trained ChIPanlayser on the top 10 regions on chromosome 3R. Top regions were selected from the 3393 regions described in *Materials and Methods*. We then validated our model on the top 20 regions on chromosome 2R and, for comparison, on top 10 regions on chromosome 3R that did not contain the training set. (**A**) shows example profiles obtained during training. (**B**) shows validation profiles obtained on chromosome 3R. (**C**) are profiles obtained during validation on chromosome 2R. Finally, (**D-G**) are the associated metrics for training and validation: AUC, Spearman correlation, recall and MSE respectively.

Finally, we compared the performance of ChIPanalyser to other TF binding prediction frameworks namely PIQ, msCENTIPEDE and Catchitt. As many available tools and frameworks are restricted to only considering human or mouse data, we selected CTCF ChIP data in astrocyte cells (*Homo sapiens*) as provided by ENCODE (Table S1 in Supplementary Materials). This data set was also used in the DREAM challenge competition related to TF binding prediction. It should be noted that neither PIQ nor msCENTIPEDE have a validation step and, for this reason, we ran both PIQ and msCENTIPEDE on both chr11 and chr18 of the human genome (full chromosomes). Input BAM files were truncated using samtools to only include these chromosomes. As PIQ and msCENTIPEDE provided discrete TF binding sites, we smoothed scores over 100 bp in order to keep the evaluation window consistent between all tools. Catchitt was trained on chr18 while ChIPanalyser was trained on the top ten regions of chr18. We then validated each tool on varying number of regions on chr11 (20, 50, 100, 200, 500, 1000 and 6755 bin of 20 Kb). ChIPanalyser outperforms all other tools when the number of regions used for validation is below 500 (see Figure 4 A-C). When using more than 500 regions for validation, we observed that all tools performed similarly poor. This trend holds true when using AUC, recall and MSE as goodness of fit metrics (although less clear with MSE). Furthermore, we trained all tools (when possible) on whole chr18 and validated on whole chr11. This ensures that all tools were trained and validated using the same data. We show that all tools perform similarly poor when using the ChIP enrichment method to estimate goodness of fit (see Figure 4D-F).

**Figure 4:**
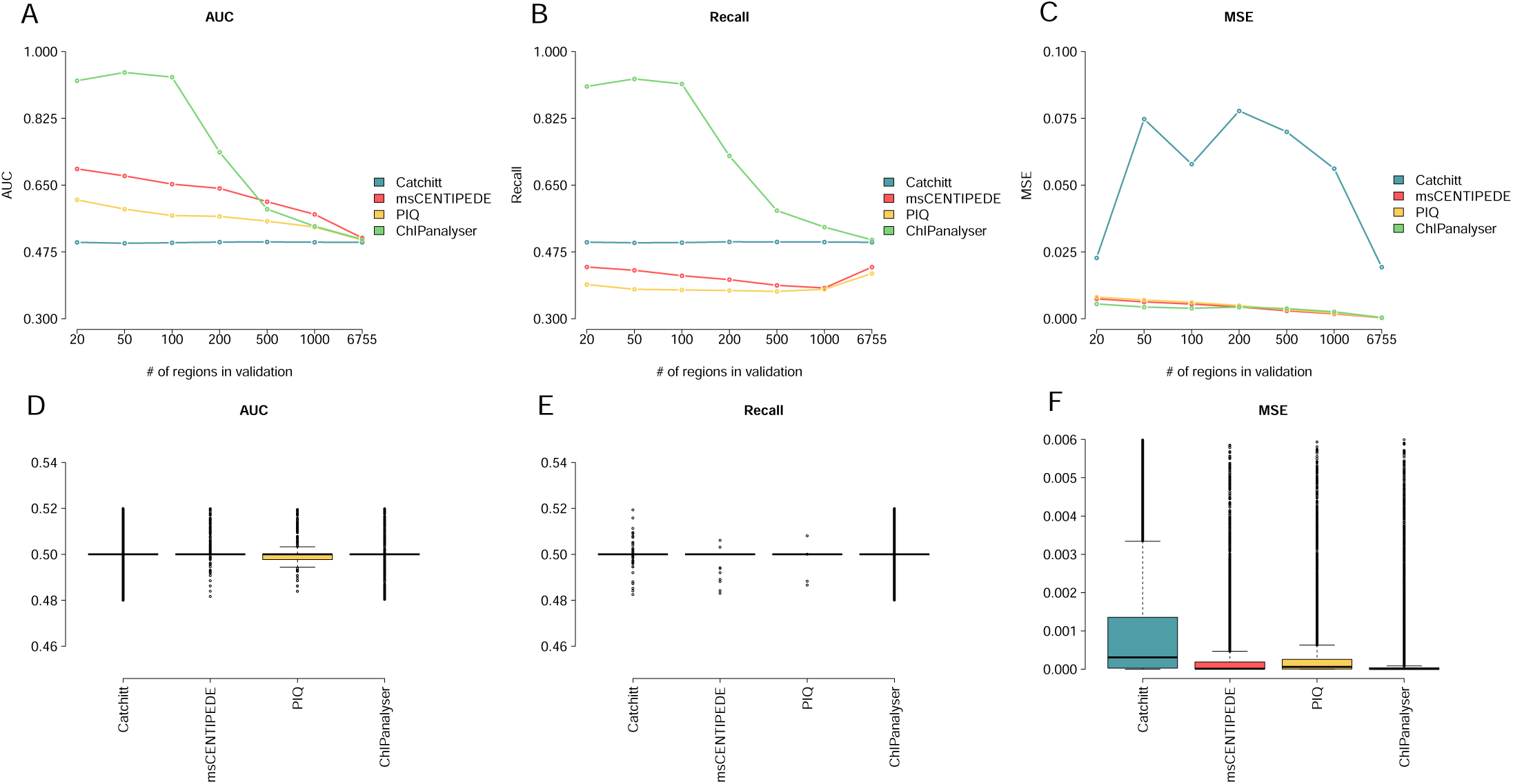
Performance Comparison between TF binding predictions tools. After training each model in their respective training set, we validated each tool using varying number of validations regions. ChIPanalyser out performs other tools when number of validation regions remains below 500. This demonstrates ChIPanalyser’s ability to describe TF binding behaviour with respect to peak strength. (**A**) shows AUC scores between Catchitt, msCENTIPEDE, PIQ, and ChIPanalyser over the selected validation regions in chr11 on *Homo sapiens*.(**B** and **C**) are respectively recall and MSE over validation regions for each tool. Finally, (**D**), (**E**), and (**F**) show the performance of all tools when trained on whole chr18 and validated on whole chr11. It should be noted that these results were performed using the ChIP enrichment method (see Methods) and that this approach considers both ChIP peak location as well as local peak enrichment.

### DNA accessibility plays a key role in the binding of TFs

Steric hindrance can influence the binding of some TFs to DNA, meaning that a TF molecule would only bind stretches of DNA if they are accessible. Any given genomic region can be considered either accessible or inaccessible and that is sufficient to explain the binding profiles of most TFs [13]. Here, we selected accessible DNA based on DNase Hypersensitivity Sites (DHS) in three *Drosophila* cell lines (Kc167, S2 and BG3). In these circumstances, DNA was either considered accessible (score of 1) or inaccessible (score of 0). As a point of comparison, we also considered all DNA to be accessible (No Access – all regions are assigned a score of 1) and also used a min-max normalised DNase score as continuous DNA accessibility level (a continuous value between 0 and 1). We focused our analysis on three TFs: CTCF, BEAF-32 and su(Hw). We trained our model on the top 10 regions for each data set. Then, we validated our results using the optimal parameters selected during training. The optimal parameters were selected by minimising MSE between experimental ChIP profiles and predicted ChIP profiles. Validation was carried out on the top 100 regions for each dataset (excluding the ones used for training). Figure 5 shows that, for BEAF-32, the binding predictions were improved when considering DNA accessibility. Nevertheless, su(Hw) and CTCF displayed a different behaviour, as the mean AUC decreased when DNA accessibility was considered for most ChIP-seq datasets (Figure 5A-B). This difference is especially striking in the case of su(Hw). The performance of the model drastically improves when all DNA was considered accessible or when we used continuous values for DNA accessibility. CTCF showed a similar trend although improvement was not as striking as in the case of su(Hw). This would indicate that only a small number of CTCF peaks are located in closed chromatin regions that display intermediary levels of accessibility.

**Figure 5:**
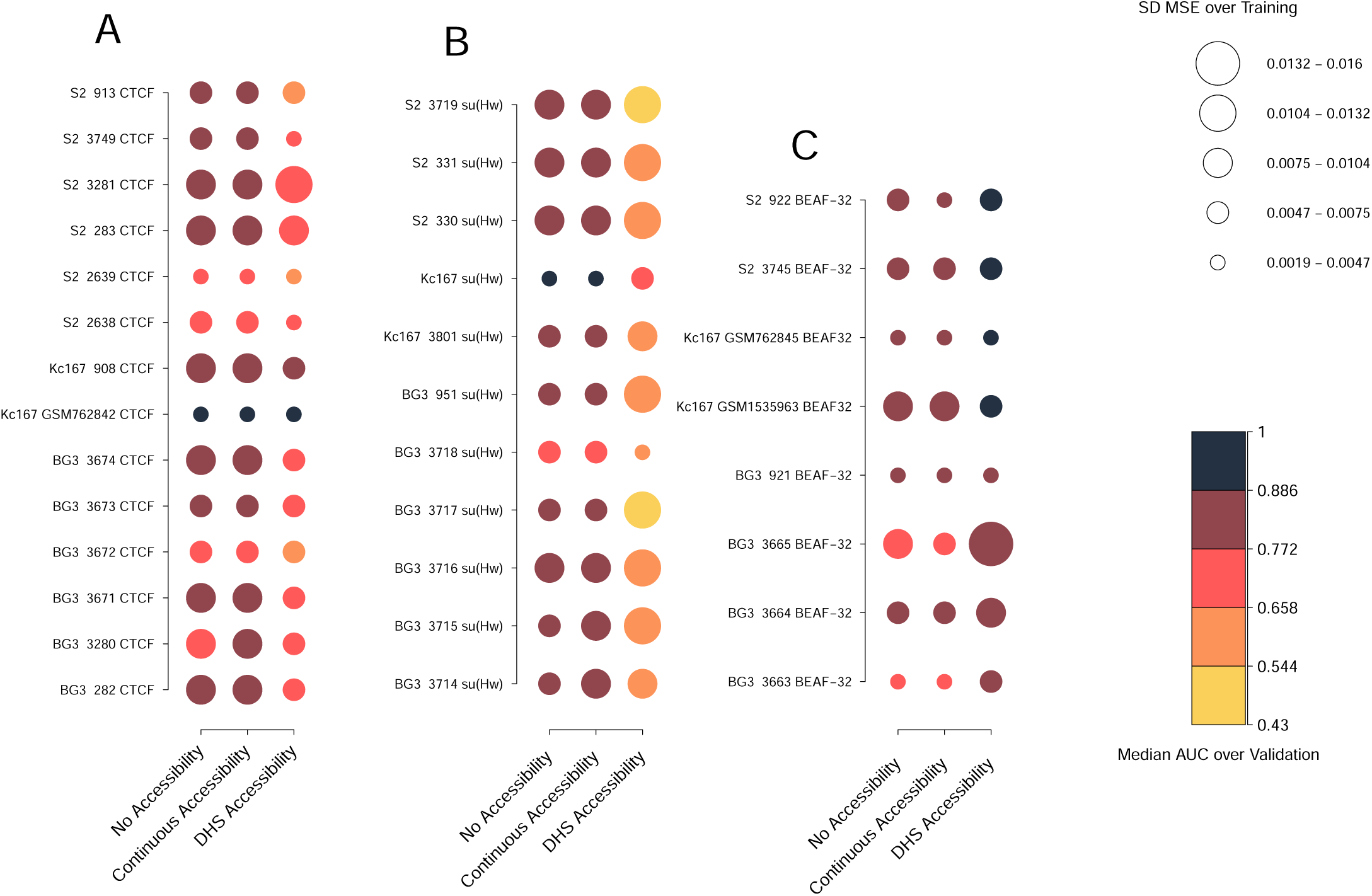
DNA accessibility, number of molecules and binding energy have different roles in TF binding. We selected optimal parameters by minimising MSE over the training set (see Table S3 in Supplementary Materials) and then computed the median AUC scores over the top 100 regions in the validation set. We considered different ChIP replicates in S2, Kc167 and BG3 cells for: (**A**) CTCF, (**B**) su(Hw) and (**C**) BEAF-32. Darker colours indicate higher AUC scores, while lighter colours lower AUC scores. We also investigated the influence of number of bound molecules and scaling factor on TF binding by computing the standard deviation of MSE scores for all combination of parameters over the training set. Smaller circles indicate less variability in MSE when different parameters are used and larger circles more variability.

While DNA accessibility seems to play a role in the quality of our predictions, we also observed that the number of bound molecules (*N*) and scaling factor (*λ*) show a reduced influence when DNA accessibility is considered for CTCF (Figure 5). In particular, we observed less variation in MSE for different sets of parameters, when DNA accessibility was included, i.e., larger circles indicate that number of bound molecules and *λ* have a more important role in TF binding, while smaller circles indicate that they have a less important role. This opposite trend is seen in the case of su(Hw) where *N* and *λ* show an increased influence when DNA accessibility is considered. BEAF-32 on the other hand is negligibly influenced by *N* and *λ* independently of whether or not we consider DNA accessibility. The rational behind this approach was that if different combinations of parameters produce strong difference in goodness of fit, then *N* and *λ* play a strong role in producing our predicted profiles. On the other hand, if we observed low variation in MSE, we could conclude that regardless of the values assigned to these parameters, the predicted profiles would remain similar.

To factor in for potential differences in the capacity of the model to predict binding in regions with strong or weak ChIP signal, we trained ChIPanalyser on the top 10 regions (see Methods) for each data set and then selected the top 20, 50, 100, 150, 200, 500, 1000 and 3283 regions for validation (excluding regions used for training). We looked at how the median AUC scores (over all data sets) changes when regions with weaker binding are included in the analysis or when DNA accessibility is considered. For each number of regions selected for validation and for each data set, we subtracted the mean AUC score when no accessibility was considered from the AUC score with DHS accessibility (Delta mean AUC). First, we observed that CTCF exhibited a slightly lower AUC score when DNA accessibility was considered (Figure S6A and D; see also Figures S7A-S10A in *Supplementary Figures*). The decrease in AUC scores observed upon considering more regions (see Figure S7A in *Supplementary Figures*) implies that CTCF binds preferentially to genome hotspots. CTCF shows strong binding at only a subset of binding sites. Interestingly, the same results were found in human data sets as described in Figure 4A-C. In contrast to CTCF, BEAF-32 displayed higher AUC scores when DNA accessibility was included, supporting the previous findings (Figure S6B and E; see also Figures S7B – S10B in *Supplementary Figures*). BEAF-32 AUC scores were not affected by the increase in the number of regions (Figures S6B and E and Figures S7B – S10B in *Supplementary Figures*), which means that BEAF-32 binding is not influenced by the number of regions selected. In other words, BEAF-32 would bind anywhere along the genome as long as it has an accessible site. In this context, we call BEAF-32 a global binder and CTCF a hotspot TF.

Furthermore, Supplementary Figure S6C and S6F shows that there is a strong and statistically significant (*p* < 0.05) reduction in AUC score for su(Hw) when DNA accessibility is included, which indicates that su(Hw) would bind in less accessible DNA (also Supplementary Figures S7C-S10C). While, su(Hw) did not generally perform well when DNA accessibility is considered, the performance of our model to predict su(Hw) binding is also tied to the number of regions selected and our results show that the strongest su(Hw) binding sites are found within inaccessible DNA. As the model uses experimental ChIP data for training, these results suggest that many su(Hw) peaks are located in inaccessible DNA.

### Number of bound molecules and TF specificity plays a limited role in the binding of architectural proteins

To investigate the robustness of our estimated parameters, we computed the optimal parameters for different biological replicates. Despite strong variations between experimental data, we show that the predicted optimal set of parameters when using MSE remained similar between biological replicates (see Supplementary Figure S11). This suggests that despite biological and technical variation between replicates performed by different labs using different protocols, our model robustly infers a similar number of bound molecules and scaling factor for a given TF.

Interestingly the importance of method selection is clearly shown when considering other metrics (see Figures S12 – S14 in *Supplementary Figures*). The optimal parameters estimated over the training set can be found in Table S3, Table S4, Table S5 and Table S6 in *Supplementary Tables* for MSE, AUC, recall and Spearman correlation coefficient.

To investigate the influence of these parameters, we assumed that a high variation of goodness of fit score for each combination of parameters would suggest a strong influence of these parameters on TF binding. If goodness of fit scores varied little between parameter combinations, we can then conclude that they do not strongly influence our predicted profiles. We then analysed the standard deviation of MSE over training between different sets of parameters and we found that some TFs are not strongly influenced by the number of bound molecules or the scaling factor (described by circle size in Figure 5).

CTCF showed a slight decrease in sensitivity to number of bound molecules and the scaling factor when accessibility was considered (Figure 5A), while, for BEAF-32, *N* and *λ* showed reduced influence on the binding profile (Figure 5C). In contrast to CTCF, su(Hw) displayed an increased sensitivity to *N* and *λ* only when DNA accessibility was considered (Figure 5B). This means that DNA accessibility would be the strongest driver towards predicting TF binding of these architectural proteins. Restricting the amount of available binding motifs would be more influential than TF copy number and the ability of a TF to discriminate between high and low affinity sites. Interestingly, this still holds in the case of su(Hw); we show that su(Hw) binding sites are most likely found in less accessible DNA. Our results suggest that relative TF abundance only play a role on binding sites found in accessible DNA.

### ChIPanalyser recapitulates TF binding profiles in different cell lines by considering relative mRNA abundance

We wanted to further investigate the predictive capabilities of our model and also demonstrate its mechanistic soundness for CTCF, BEAF-32 and su(Hw) in the three selected cell lines. For that, we estimated the optimal set of parameters in one cell line and aimed to predict TF binding in a different cell line taking into account changes in DNA accessibility using DHS data and changes in number of bound molecules using relative changes in RNA abundance. For example, we estimated the optimal set of parameters for CTCF in Kc167 cells that would minimise MSE as *λ* = 1.5 and *N* = 10^4^ over the top 10 regions (see Methods). By rescaling *N* based on relative RNA-seq levels of CTCF in the two cell lines, we could approximate the number of CTCF molecules bound to DNA in BG3 cells (*N* ≈ 1.6 × 10^4^). This together with BG3-specific DNA accessibility data is capable of predicting the ChIP-seq profile in BG3 cells (see Figure 6A and B). RNA rescaling of the number of bound molecules seems to recover both the number of peaks and their location with high accuracy. The rescaling of number of bound molecules lead to differences in terms of MSE variation between estimated and rescaled (Figure 6G).

**Figure 6:**
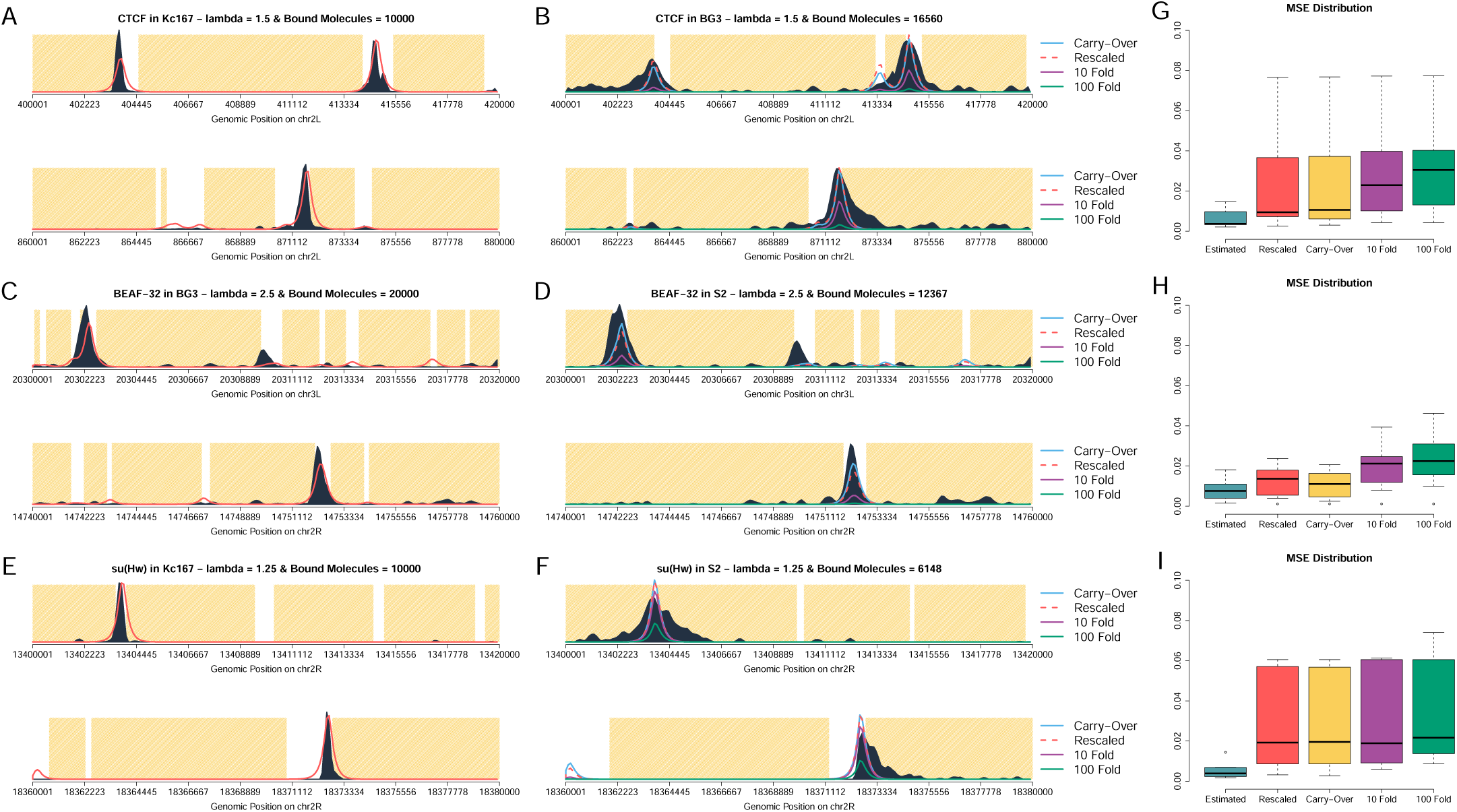
TF abundance remains stable between different cell lines when considering relative mRNA abundance. **A-F** show predicted ChIP-seq profiles with our TF abundance estimated based on RNA-seq. The yellow area represents inaccessible DNA, the dark area represents experimental ChIP signal and the red lines are our predicted profiles. We estimated the number of bound molecules in one cell line (**A, C** and **E**) and rescaled our estimate using relative mRNA abundance in an other cell line (**B, D** and **F**). (**B, D** and **F**) The dashed red line represents the rescaled value of number of bound molecules based on relative RNA-seq abundance, the light blue the original value estimated in (**A, C** and **E**).The purple line and the green line represent the original estimated value reduced 10 and 100 times respectively. (**G**, **H** and **I**) Boxplots with MSE for all cases in the estimated and predicted profiles at top 10 regions for both training and validation.

The estimated MSE (MSE over the training set) in one cell line is lower than its counter parts in the other cell line. However, we attribute this difference to differences in data quality between cell lines. This is especially striking in Figure6 A, B, E, and F. ChIP peaks in the training cell line (see Figure6 A and E) display shaper peaks and much less background signal then ChIP peaks in the validation cell line (see Figure6 B and F). As described in Methods, ChIPanalyser estimates goodness of fit using ChIP enrichment scores and therefore is sensitive to background signal and/or wider than expected peaks. The same analysis was performed for BEAF-32 (Figure 6C, D and H), where we estimated parameters in BG3 cells (*λ* = 2.5 and *N* = 2 × 10^4^) and rescaled the number of molecules in S2 cells (*N* ≈ 1.2 × 10^4^). Once again, the model correctly predicts ChIP profiles in both location and relative enrichment. Finally, for su(Hw) (Figure 6E, F and I) we estimated parameters in Kc167 cells (*λ* = 1.25 and *N* = 10^4^) and rescaled the number of molecules in S2 cells (*N* ≈ 6 × 10^3^). Again, the predictions of the model are accurate.

Our results show that ChIPanalyser can accurately recapitulate ChIP profiles between cell lines using cell specific DNA accessibility data and number of bound molecules. Nevertheless, we still do not know which of the two is the more important factor or whether both have similar contributions. To address this, we also assumed that in the predicted profile that there is *(i)* no change (same number of bound molecules is used in both cells), *(ii)* a 10 fold reduction and *(iii)* one 100 fold reduction in the number of bound molecules and repeated the analysis. Figure 6 shows that using the same TF abundance as in the original cell line produces extremely similar ChIP like profiles. In fact, we observed a significant reduction in the predicted profile only when reducing the number of bound molecules by 100 (for CTCF and su(Hw)) or 10 (for BEAF-32) fold. These results show that cell differences in binding profiles of TFs, at their *strong binding regions*, would mainly come from differences in DNA accessibility and not relatively small changes in TF abundance. The only way that TF abundance could impact the binding profile (and, consequently, lead to changes in gene regulation) is when the expression of the TF is strongly down-regulated.

### Hox genes show differential binding preferences with respect to DNA accessibility

Hox proteins are key players during development. Recently it has been suggested that Hox proteins show different binding preferences with respect to DNA accessibility [33]. Most notably, Ubx and Abd-A would bind predominately in open chromatin, while other Hox TF (Lab, Pg, Dfd, Scr and Abd-B) would prefer closed chromatin. We selected three Hox TFs (Ubx, Dfd and Abd-B) and ran our model using different levels of DNA accessibility. DNA accessibility levels were selected based on quantile distribution of ATAC-seq scores (see Methods). This means that higher QDA scores lead to fewer regions being marked as accessible.

We trained our model on the top ten regions selected from the 3838 selected for the Hox analysis (see Methods) for each QDA accessibility. Our results show that Ubx exhibits a preference towards open chromatin. In Figure 7A, the maximum AUC score for Ubx increases with the increase of the QDA score. Dfd and Abd-B on the other hand were not strongly influenced by QDA accessibility. This means that these TFs can bind in inaccessible DNA. According to our model, Ubx performed best with 0.99 QDA (top 1% ATAC-seq scores – AUC 0.928), while Abd-B and Dfd with 0.95 QDA (top 5% ATAC-seq scores) and 0.8 QDA (top 20% ATAC-seq scores) respectively (see Figure 7B). It should be noted that these scores are on the training set as the goal was to understand how QDA would effect the training of our model. We then validated our model on the top 100 regions (excluding the ones used for training) using the optimal set of parameters inferred during training and plotted the predicted profiles for Hox TF (see Figure 7C,D, and E).

**Figure 7:**
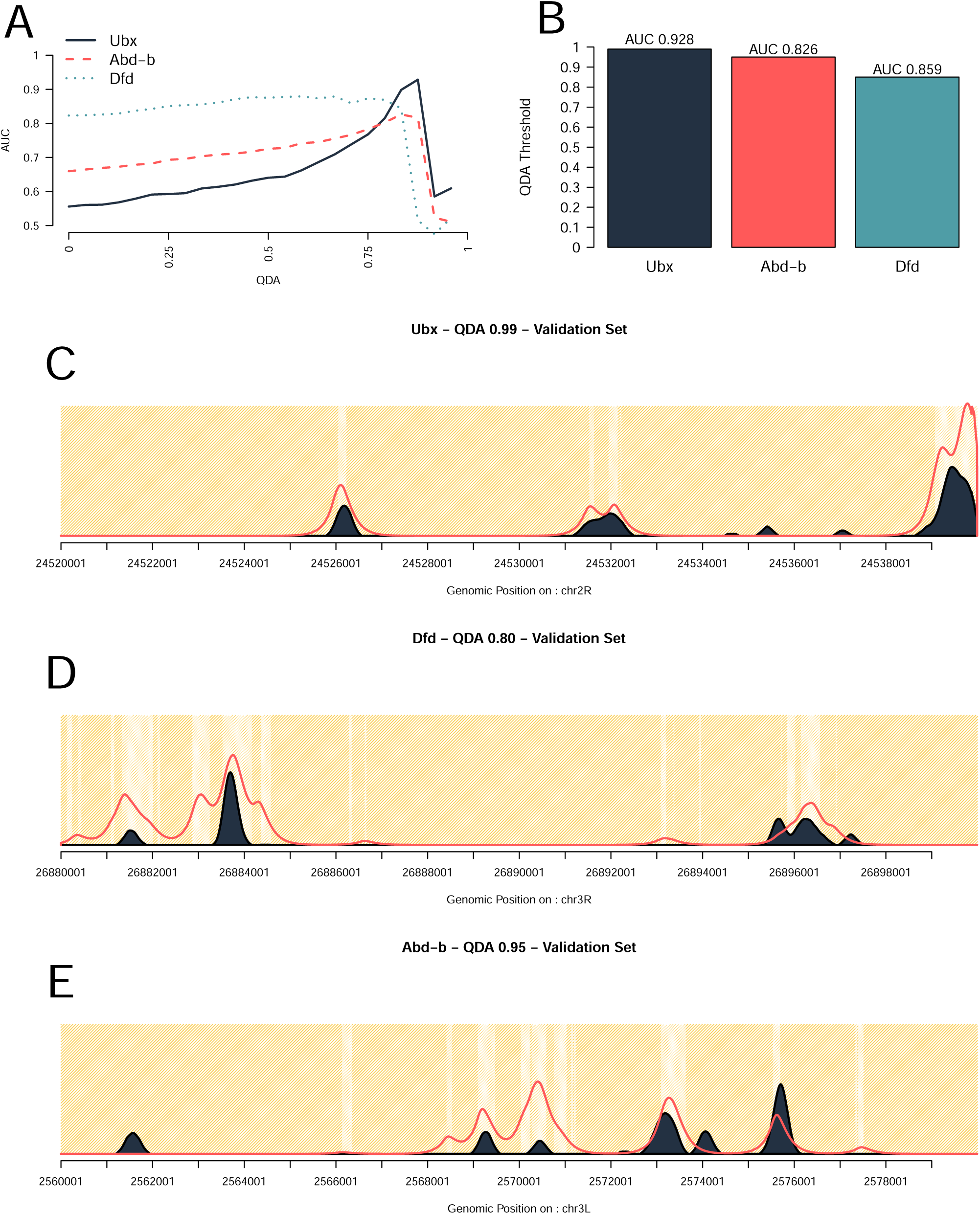
Hox genes show binding preferences towards DNA accessibility. We tested our model using different DNA accessibility stringencies. (**A**) Maximum AUC score as a function of stringency of DNA accessibility (the higher the QDA value the less DNA is called accessible) for three Hox TFs: Ubx, Dfd and Abd-B. (**B**) The best performing QDA accessibility in terms of AUC. (**C, D** and **E**) Binding profiles and prediction of the ChIP data at individual *loci* taken from the validation set for the three TFs.

The model recovers the position of peaks accurately especially for Ubx (see Figure 7C-E). While for Dfd and Abd-B most of the peaks are detected, their height is not always an accurate representation of the strength of the ChIP-seq signal. Hox TFs are known to display cooperative interactions and there are reports that both Dfd and Abd-B have a higher number of sites in the bound peaks, suggesting they bind cooperatively to open the chromatin [33]. Our model does not include cooperative interactions and this could explain the reduced performance for Dfd and Abd-B.

## DISCUSSION

Our analysis shows that ChIPanalyser and its underlying model predicts binding profiles of TFs (ChIP) with high accuracy and most importantly it can also shed light on the binding mechanism of TFs. We show how ChIPanalyser not only predicts location of peaks, but can correctly predict the enrichment of a TF at a given location.

### TFs used different binding mechanisms

In this analysis, we focused our attention on three DNA binding proteins: CTCF, BEAF-32 and su(Hw). All three TFs are known architectural proteins in *Drosophila* but also play roles in transcription regulation and insulation [35, 36]. Moreover, it was shown that these three TFs have distinct binding behaviours and were classified into three subclasses with respect to chromatin architecture [37, 38]. In our analysis we show that they all exhibit different behaviours with respect to DNA binding.

Our findings suggest that CTCF binds to hotspots along the genome and this could be explained by the observation that the strongest peaks are in fact highly conserved binding sites. CTCF binding to highly conserved sites can be explained by our model, but something else is responsible for the reduced binding at less conserved sites (i.e. cell specific CTCF binding) [39].

BEAF-32 is a *Drosophila* specific insulator [40] that shows preferential binding towards TAD boundaries, but also is involved in transcription itself [41]. Previous studies showed that BEAF-32 has uniform binding along the entire genome [37]. Our results confirm that BEAF-32 shows a strong preference towards accessible DNA and that the majority of accessible sites would be bound.

Furthermore, we show that su(Hw) binds in both open and closed chromatin. su(Hw) plays a role in chromatin insulation and remodelling [42] and is also a primary actor in the interaction between the genome and nuclear lamina [43]. This would explain why su(Hw) can bind in both open and closed chromatin and why ChIP peaks might not overlap well with DNase hypersensitivity data. It has also been shown that su(Hw) binding sites tend to cluster together (with varying number of sites) and that these sites are constitutively bound by su(Hw) [44]. Interestingly, it seems that only isolated high affinity sites had a role in transcriptional regulation and the clustered sites were more involved in chromatin architecture.

### DNA accessibility is the main driver of binding to DNA for some TFs

Our results show that DNA accessibility and number of bound molecules control the binding profiles of TFs (Figures 5). When we estimated the binding parameters (*λ* and *N*) in one cell line and then predicted TF binding profiles in a different cell line based on changes in DNA accessibility and number of TF molecules (using changes in mRNA), we found a good agreement between our predictions and the actual ChIP-seq dataset (see Figure 6). Nevertheless, the changes in number of TF molecules between the two cell lines did not seem to make any difference to the predicted profiles (compare blue and dashed red line in Figure 6 B, D and F). This means that biologically relevant fluctuations in TF numbers between different cell lines would have little effect on the differences in binding profiles of TFs, which would be mainly driven by changes in DNA accessibility. Furthermore, only very strong knock-downs would decrease or deplete ChIP peaks. It should be noted that CTCF, BEAF-32 and su(Hw) are highly expressed architectural and insulator proteins and, thus, they would be expected to saturate their binding sites. Interestingly, only strong depletion of CTCF in mammalian cells (using Auxin Inducible Degradation) was able to lead to noticeable changes in 3D chromatin chromatin loops controlled by CTCF [45].

Why would changes in concentration of the TF have such a limited effect on their binding? One potential explanation is that these TFs control the expression of essential genes that should be tightly regulated to buffer fluctuations in number of molecules that affect the cell [46].

Finally, we also investigate the capacity of our model to differentiate between TFs that can bind only in open chromatin or also partially opened chromatin. Our results showed that while Ubx displays a strong sensitivity to open chromatin and binds in the top 1% accessible sites, the binding of Abd-B and Dfd is less influenced by DNA accessibility (with Abd-B and Dfd binding in top 5% and 20% respectively accessible regions); see Figure 7. Hox TFs are known for having a similar motif, but display differences in their binding profiles [47]. It was hypothesised that binding cooperativity could explain the difference in binding profiles coupled with protein sequence changes [48]. Here, we showed that differential capacity to bind in dense chromatin could also be responsible for the difference in binding profiles of Hox TFs (see Figure 7).

### Background noise and experimental artefacts remain a challenge in TF binding predictions

We found that many ChIP datasets suffer from significant background noise that would reduce our ability to accurately assess the goodness of fit of the model. Despite our approaches to reduce background noise, it seems that ChIP data will always suffer from unspecific DNA pull-down [49].

Another possibility is that the noise in ChIP signal could be the result of unspecific binding of TFs to DNA followed by one-dimensional random walk along the genome [50, 51]. Nevertheless, the washing steps in the ChIP protocol would remove this non-specific binding from the final ChIP signal [4].

We showed that choosing a goodness of fit method is context dependent. Interestingly, similarity methods (such as correlation, F-score or AUC) had the tendency to correctly call peak location but greatly underestimate the enrichment on the peak (see Figure 2). This behaviour results from the fact that these methods are highly penalised by false positive hits. The scaling factor can be described as how well a TF discriminates between a strong binding site over a weaker one. High values for the scaling factor translate to poorer ability for the TFs to discriminate between high and low affinity sites, which leads both to a higher number of false positive peaks and the model picking up smaller peaks. The number of bound molecules on the other hand, tend to affect the height of the peak (relative local enrichment). Similarity methods would avoid high values for *N* and *λ* as this would penalise their goodness of fit score more severely as opposed to dissimilarity methods (see Figure 2).

Choosing the right method will depend on the question at hand and similarity methods could be used to determine peak location, while dissimilarity metrics would be more appropriate to investigate the TF local enrichment.

### ChIP enrichment scores provide a highly stringent method to assess model performance

When comparing ChIPanalyser to other frameworks, we observed that all tools performed poorly when trained on a full chromosome (hg38 – chr18) and validated on a full chromosome (hg38 – chr11). When ChIPanalyser was trained on the top 10 regions of chr18 and validated on varying number of regions in chr11, it outperforms other tools and frameworks as long as the number of validation regions did not exceed 500.

ChIPanalyser evaluates goodness of fit using ChIP enrichment scores (see Methods). This ensures that the model considers peak enrichment during the optimisation step (both location and height of the peak). Competing tools generally assess model performance by overlapping predicted TF binding sites with ChIP peaks and do not explicitly account for peak enrichment (they assume that there is very little difference between a strong and a weak ChIP peak). While we recognise that our scoring method is best suited for TF binding events described both by peak location and peak enrichment, we selected this approach as ChIPanalyser describes a mechanistic interpretation of TF binding. ChIPanalyser was trained on the top 10 regions in order to ensure a balance between True Positive and True Negative signals leading to a more effective parameter inference. Many regions along the genome might not contain any ChIP signal and this lack of signal will affect the profiles produced by ChIPanalyser.

Finally, the goal of ChIPanalyser is not only to predict TF binding events but also shed light on the mechanisms driving TF binding. In the case of CTCF in human astrocytes (as used in the comparison with other tools), ChIPanalyser showed a decay in performance after 500 regions used for validation (see Figure 4 A-C). PIQ, msCentipede and Catchitt did not display such a clear behaviour. Interestingly, CTCF’s behaviour with respect to number of regions used for validation was also observed in our analysis in *Drosophila* data sets. These results suggest that CTCF’s binding to highly conserved sites holds true through different organisms and that ChIPanalyser was able to recapitulate this behaviour.

## Conclusion

ChIPanalyser is a user-friendly R package available on Bioconductor for predicting the binding of Transcription Factors to DNA. The package performs similarly if not better than competing tools and frameworks. More importantly, the model also provides an insight into the binding mechanisms of various DNA binding proteins. We show the nuanced role of DNA accessibility in the binding of three architectural proteins CTCF, BEAF-32 and su(Hw) in *Drosophila*. Furthermore, we demonstrate that architectural proteins are robust to relative changes in protein abundance. Finally, we recover the binding preferences of Hox TFs with respect to chromatin compaction. ChIPanlyser provides both predictive and biological modelling capabilities.

## Competing interests

Conflict of interest statement. None declared

## Funding

This work was supported by University of Essex and by the Wellcome Trust grant [202012/Z/16/Z].

## Acknowledgements

We thank Dr Rob White, Prof Sarah Bray and Zabet lab (Keerthi Chathoth, Jareth Wolfe, Romana Pop and Olivia Grant) for useful discussion and comments on the project and the manuscript. We would also like to thank Dr. Gorrie-Stone for his comments and suggestions during the development of ChIPanalyser. The analysis was performed on the HPC at University of Essex and we would like to thank Stuart Newman for his support

## Additional Files

### Supplementary Figures

Supplementary Figures as described in the main text. Figures are numbered independently of main figure numbering. All supplementary Figure are available in pdf format.

**Table 1:** Tool and framework comparison. We provide a breakdown of a few notable tools and frameworks for TF binding prediction.

|  | **ChIPAnalyser** | **Catchitt** | **FactorNet** | **Anchor** | **PIQ** | **msCENTIPEDE** |
| --- | --- | --- | --- | --- | --- | --- |
| **Language** | R | java | python 2.7 | python3.6/perl 5.1 | R | python 3.6 |
| **Organisms** | All* | All | Human | Human | All* | Human |
| **Training & Validation** | Yes | Yes | Yes | Yes | No | No |
| **Plotting** | Yes | No | No | No | No | Limited |
| **Skill Level** | Low | Intermediate | High | High | Low | Intermediate |
| **Availability** | Bioconductor | GitHub | GitHub | GitHub | bitbucket | GitHub |

### Supplementary Tables

Supplementary Tables as described in the main text. Tables are numbered independently of main Tables numbering. All supplementary Tables are available in pdf format.

## Notes

### Competing Interest Statement

The authors have declared no competing interest.

### Summary of Updates

we introduced a comparison with other tools, and new validation strategies (e.g., chromosome withhold)

