## Supplemental Figure for "Dissecting the binding mechanisms of transcription factors to DNA using a statistical thermodynamics framework"

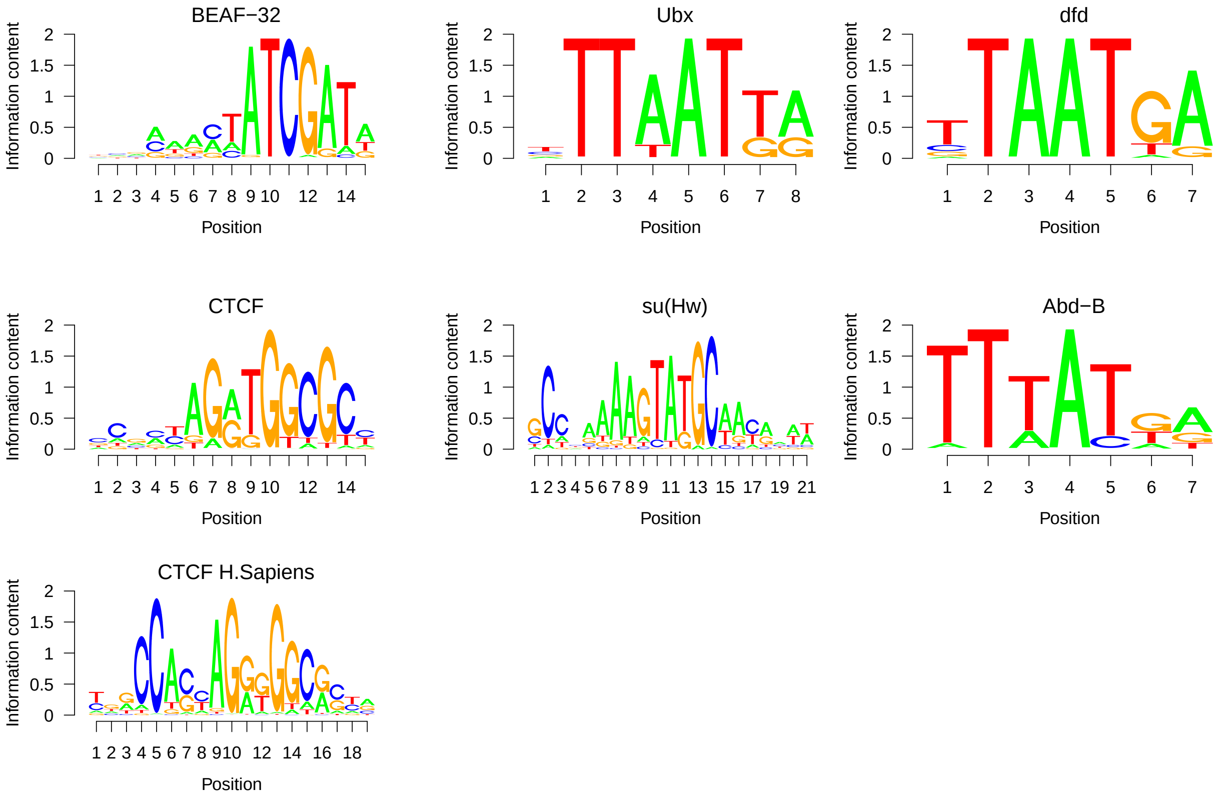


Figure S1: **Logo Motifs for all TF used in this manuscript.** Motifs were either extracted from the JASPAR database or from the MotifDB R package. In the latter case, we only considered motifs associated with the JASPAR data base. CTCF motif for *Homo sapiens* was extracted from the JASPAR database.

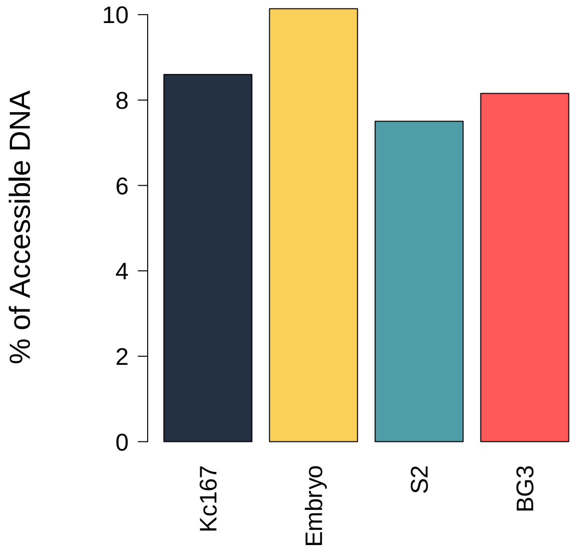


Figure S2: **Accessibility differs between cell lines.** Accessibility for three cell lines was estimated from DNase Hypersensitivity Sites (DHS) by extracting DHS broad peaks (see *Materials and Methods*). Kc167 cells displayed a slightly higher proportion of accessible DNA compared to S2 and BG3 cells. The proportions of accessible DNA remain similar to the proportion of accessible DNA in *Drosophila* embryos using the method proposed by (18).

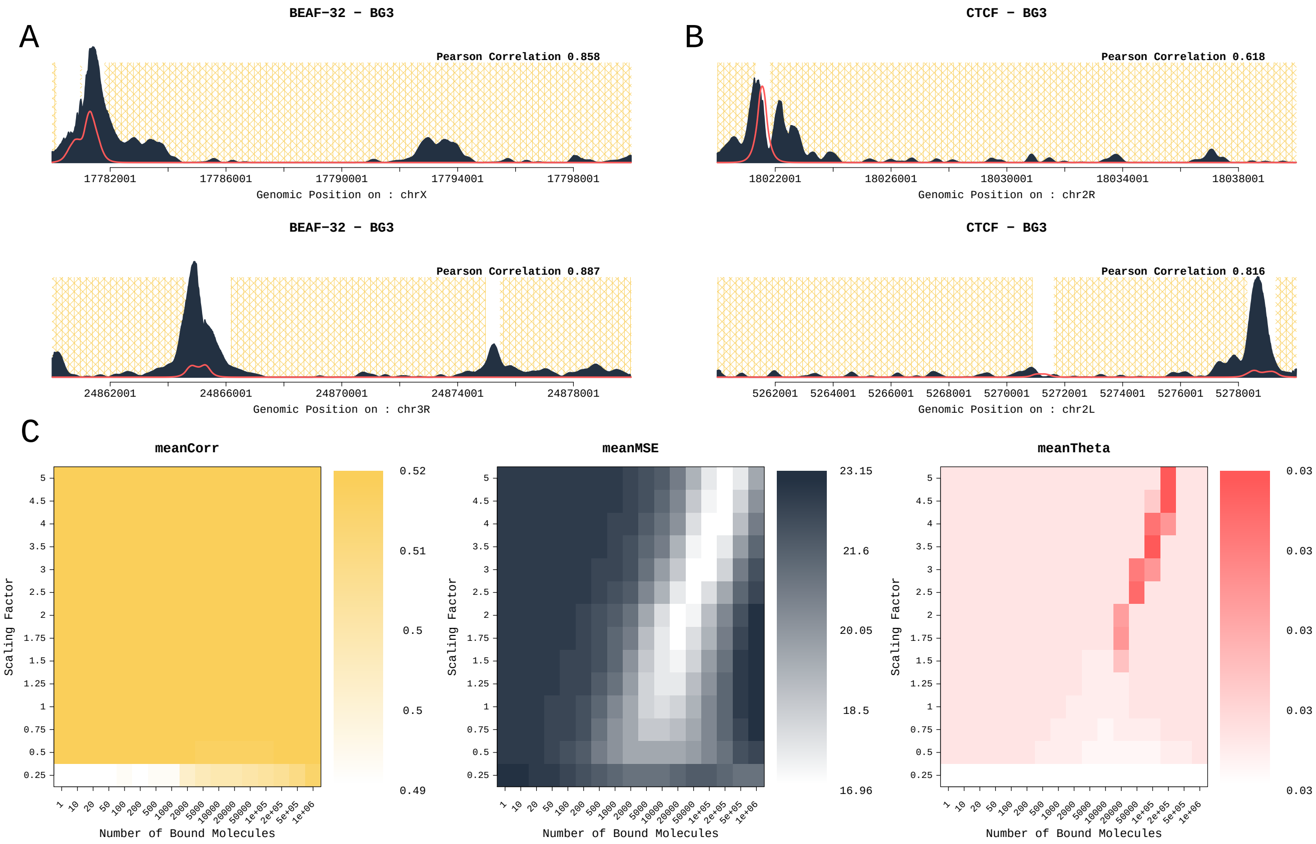


Figure S3: **The original model called for the maximisation of Pearson correlation coefficient and the minimisation of Mean Squared Error.** When extending our analysis from the five original TF, we noticed that there were certain oddities arising from the use of Pearson correlation coefficient. As seen in (**A**) and (**B**) Pearson correlation coefficient does not seem to accurately measure the goodness of fit of the model. We observed an over or underestimation of the models ability to predict TF binding. The second difficulty arising from using Pearson correlation coefficient was that it showed very little difference in correlation for some data sets. Previously, we described the optimal set of parameters by using the ratio of correlation over MSE as shown in (**C**) - meanTheta. In many cases, MSE was the driving force behind the optimal parameters. Correlation played little to no role. For these reasons, we decided to investigate other goodness of fit methods. In (**A**) and (**B**), yellow shaded areas are regions of inaccessible DNA. Dark grey is experimental ChIP signal and finally the red line represents our predicted profile. Profiles in (**A**) originate from BEAF-32 in BG3 cells. Profiles in (**B**) originate from CTCF in BG3 cells. All heat maps represent Abd-B in Kc167 cells.

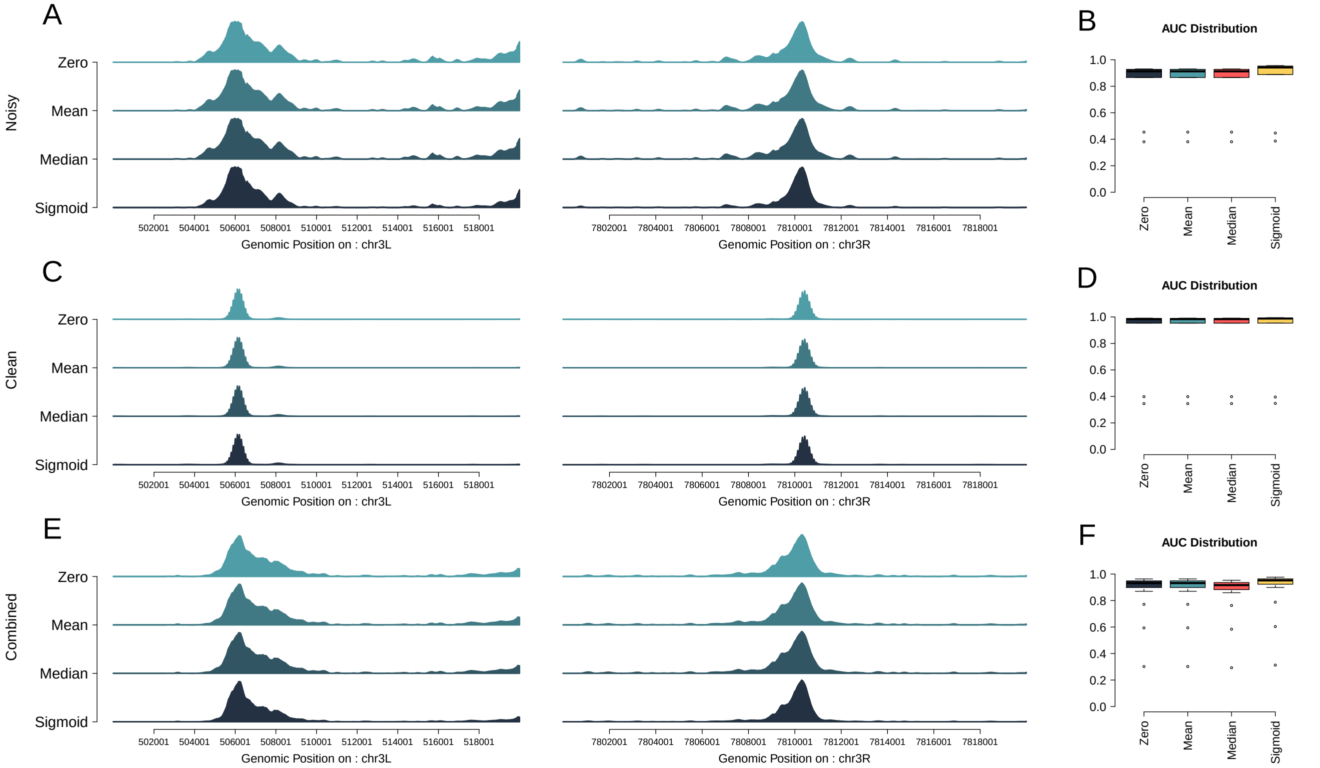


Figure S4: **Noise Filtering methods have little effect on experimental ChIP signal.** In order to improve our predictions, we sought to test four noise filtering methods: *Zero, Mean, Median* and *Sigmoid*. We tested these methods on three CTCF datasets: (**A-B**) a ChIP-seq dataset with high background noise (modEncode 3674), (**C-D**) a ChIP-chip dataset with very little background noise (modEncode 2639), and (**E-F**) a combination of all ChIP-seq datasets in S2 cells (by adding enrichment signals together at a base pair level); see Table S1 in *Appendix*. In (**A**),(**C**), and (**E**), we notice that our noise filtering methods have a limited effect on reducing noise. The consequent effect on our predictions was limited but showed a slight improvement when using the sigmoid method as described in (**B**), (**D**), and (**F**).

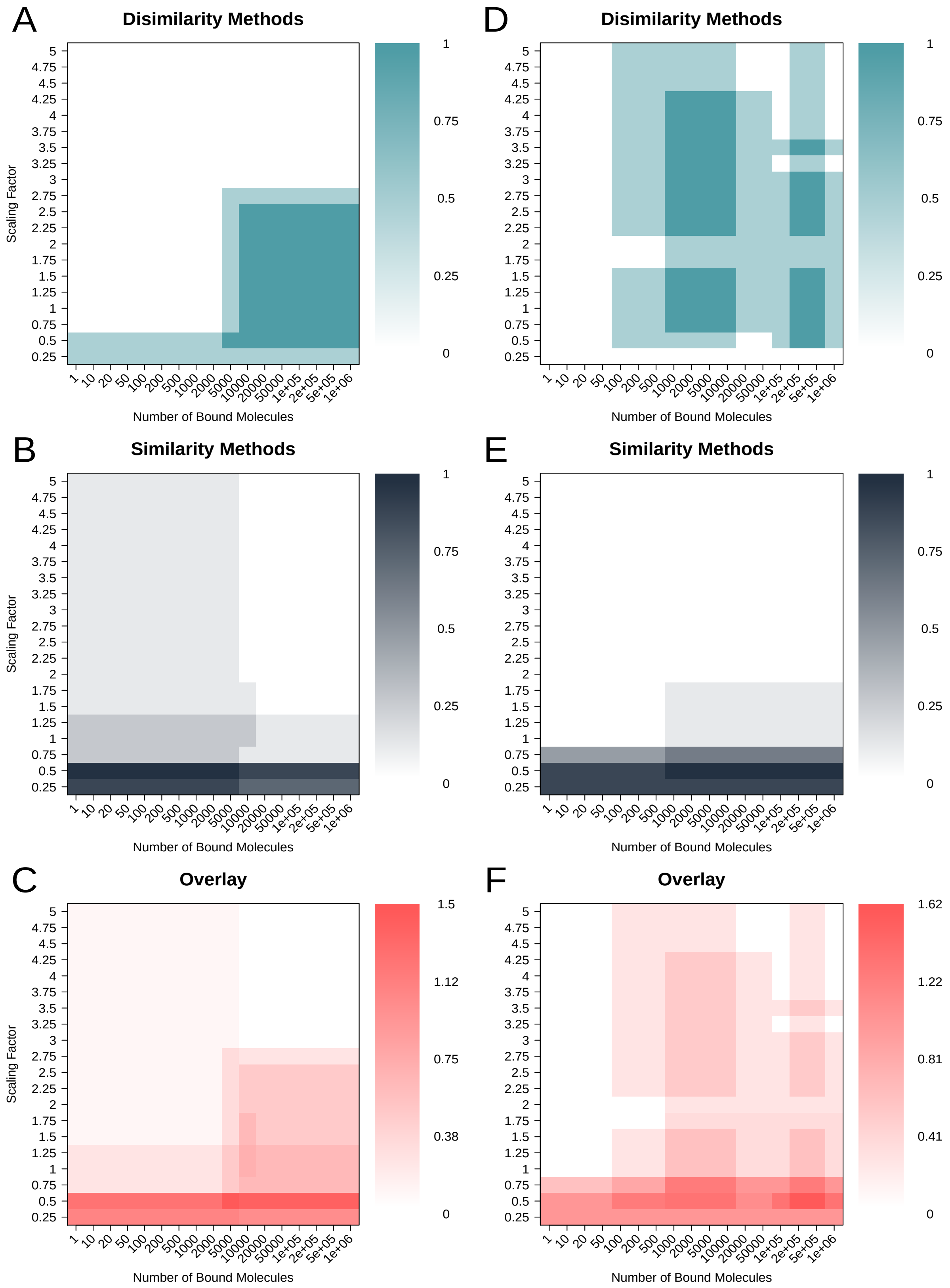


Figure S5: **Goodness of fit metrics influences optimal parameters selection** Heatmaps show the overlap of best performing (top 10 %) combination of parameters for similarity and dissimilarity methods as well as an overlay of all methods. We produced these heatmaps using the noisy (**D-F** - modEncode 3674) and clean (**A-C** - modEncode 2639) data sets.

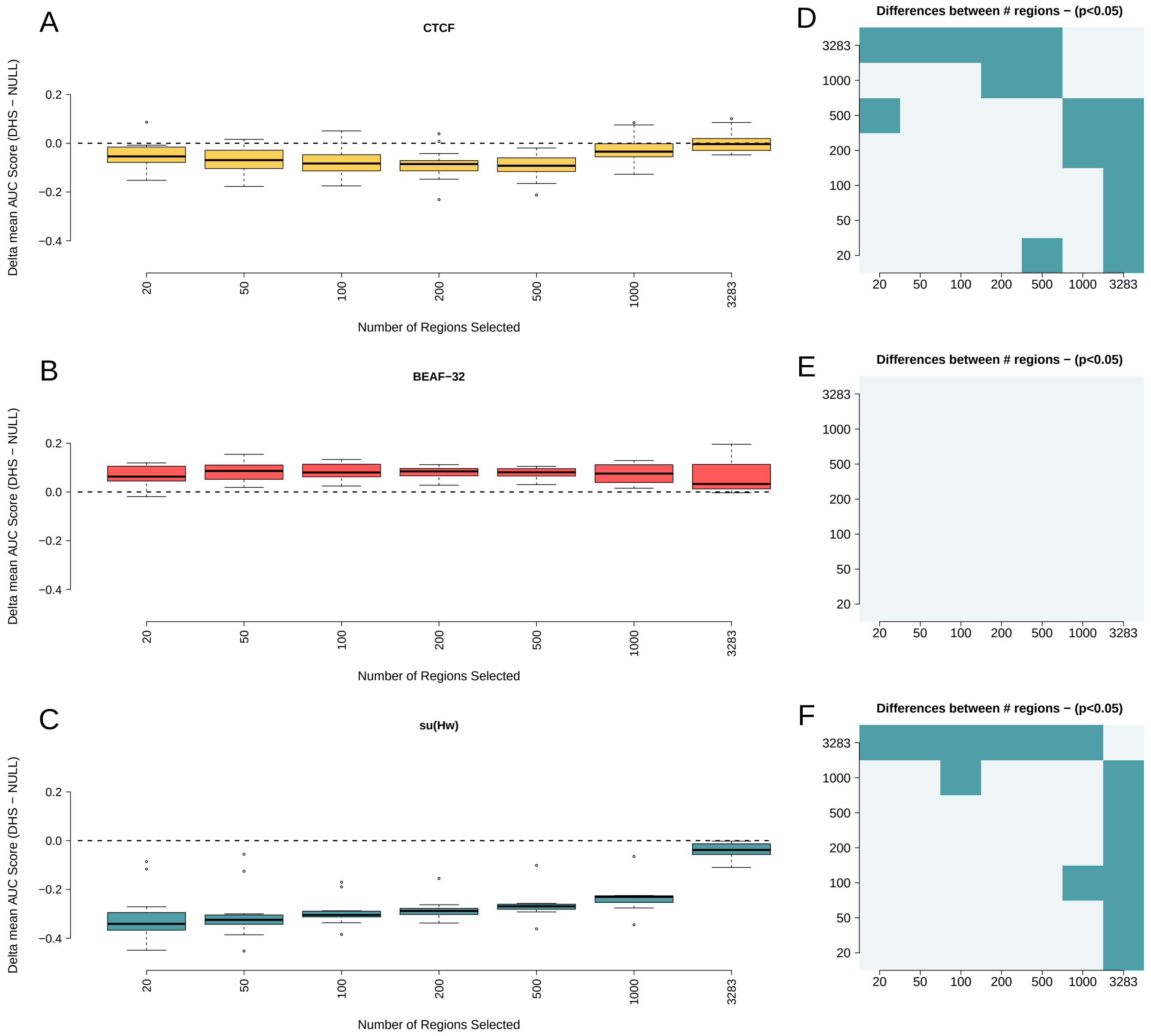


Figure S6: **Number of selected regions sheds light on TF behaviour.** (**A-C**) Boxplot representing the difference in AUC (over validation sets) between the model with and without DNA accessibility for several biological replicates and different number of selected bins. (**D-F**) T-test to assess whether the differences are statistically significant (blue indicates statistically significant differences, while light grey represent non significant combinations). (**A** and **D**). Predictions of CTCF binding shows CTCF's ability to bind to less accessible DNA. The effect of DNA accessibility decreases as the number of regions used for selection increases. Predictions of BEAF-32 binding are improved by DNA accessibility and are not affected by number of regions selected.(**B** and **E**). su(Hw) performs better when all DNA is considered accessible (**C** and **D**)

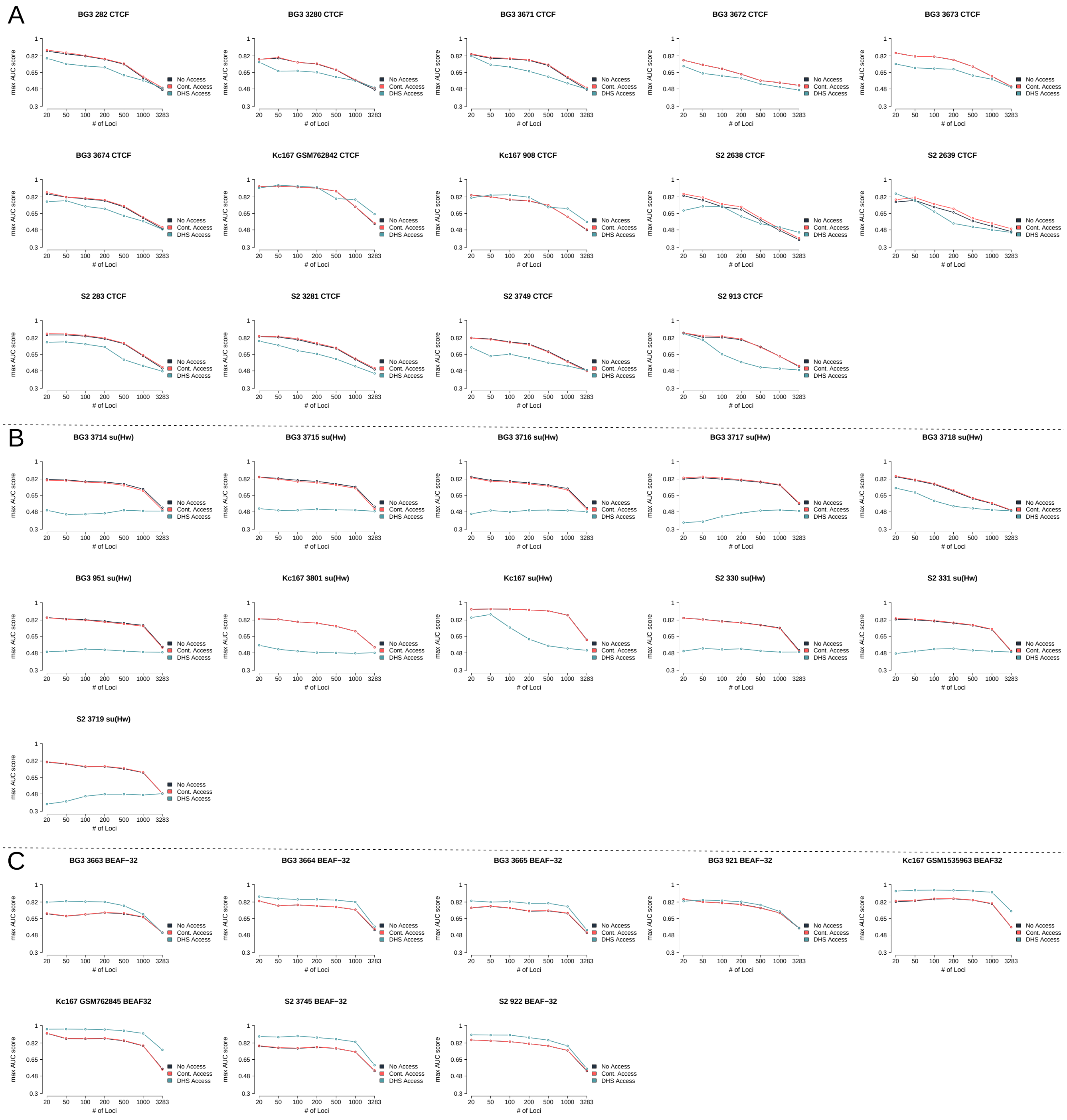


Figure S7: **Changes in AUC as the number of regions used for validation increases** We selected varying numbers of regions to validate the model. Here we present the AUC values for every CTCF, BEAF-32 and su(Hw) data set used in this manuscript; see Table S1 in *Appendix*. For each individual plot, we considered the cases of No accessibility, Continuous accessibility and DHS accessibility.

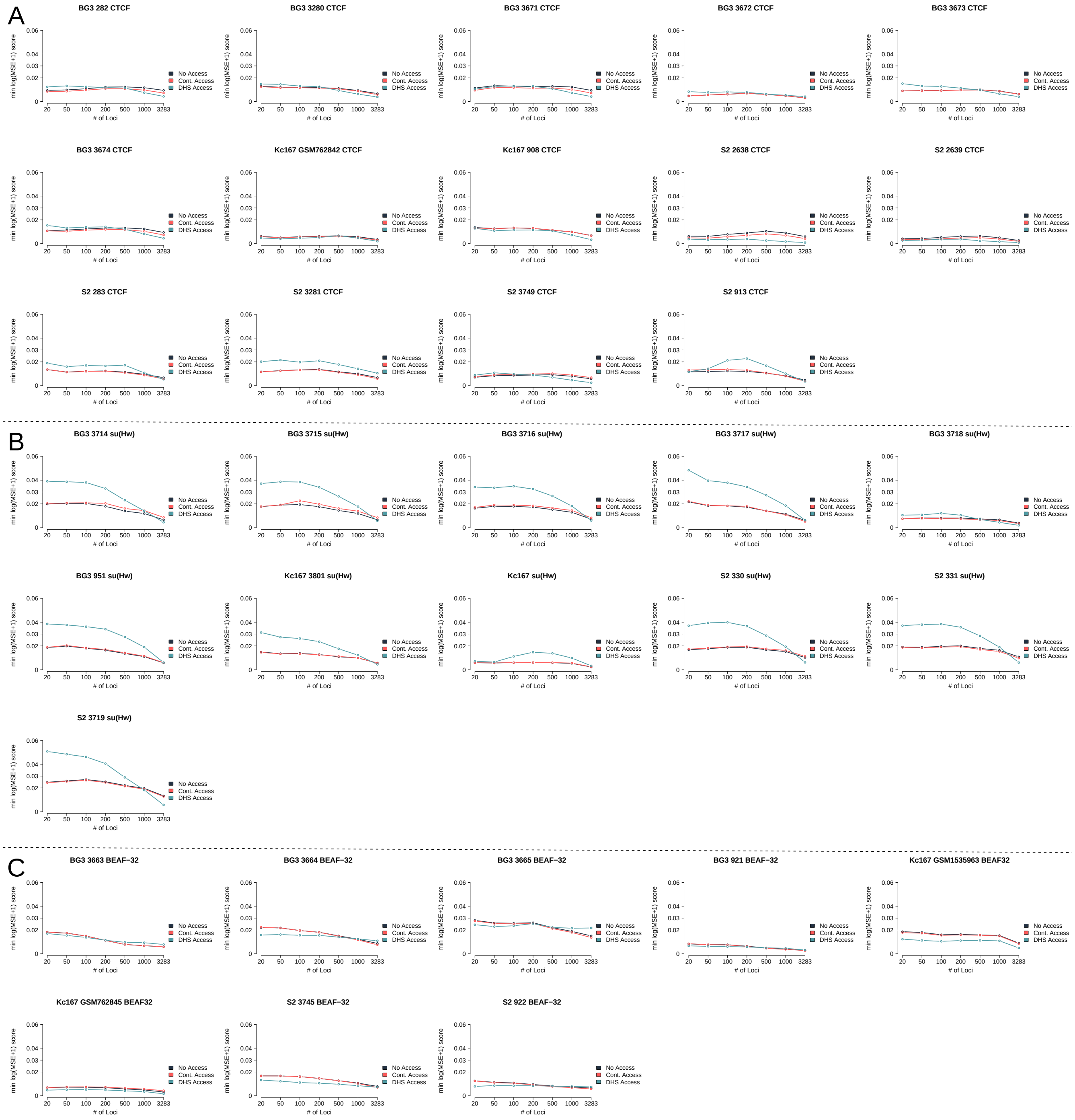


Figure S8: **Changes in MSE as the number of regions used for validation increases** We selected varying numbers of regions to validate the model. Here we present the MSE values for every CTCF, BEAF-32 and su(Hw) data set used in this manuscript; see Table S1 in *Appendix*. For each individual plot, we considered the cases of No accessibility, Continuous accessibility and DHS accessibility.

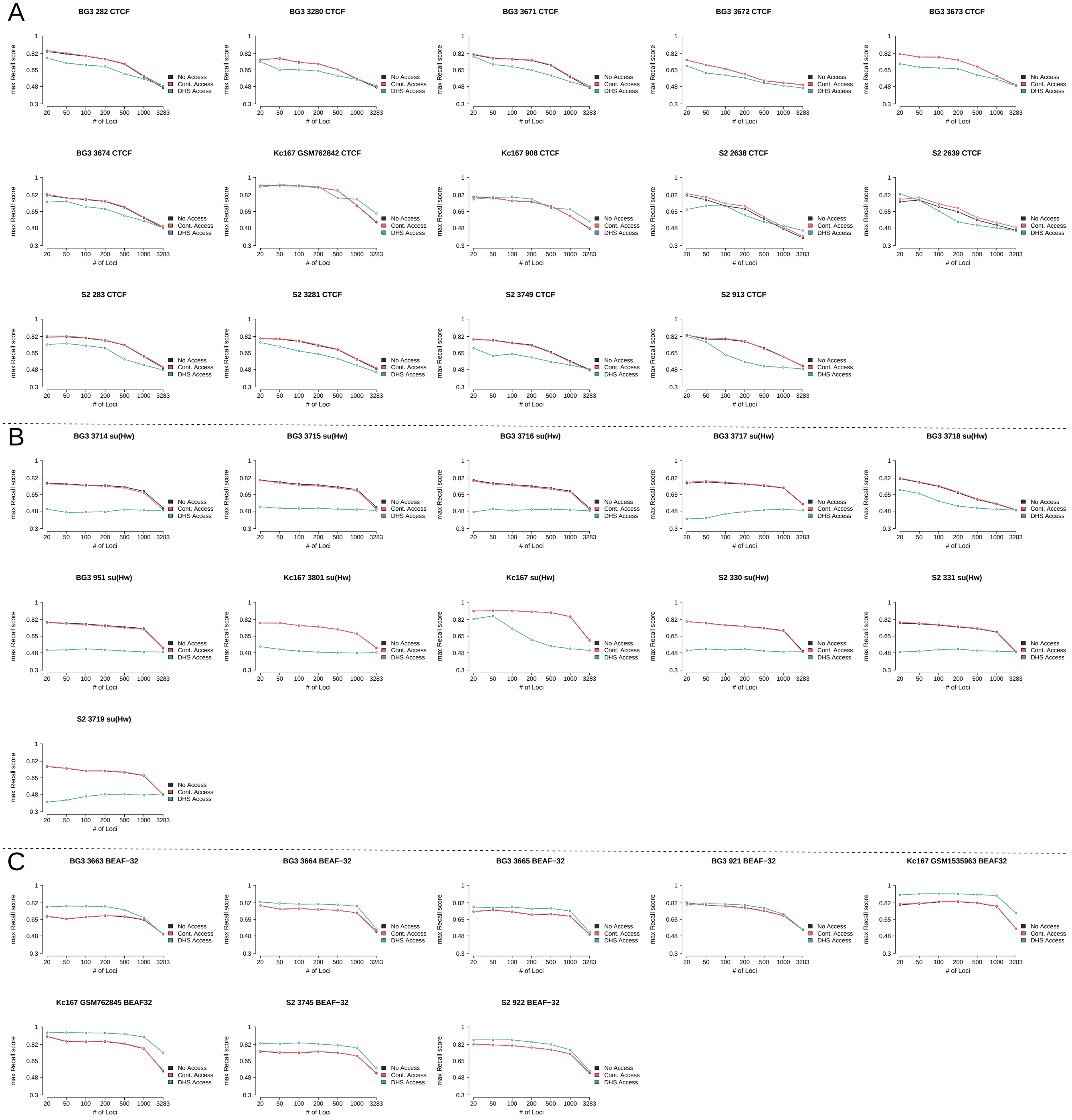


Figure S9: **Changes in recall as the number of regions used for validation increases** We selected varying numbers of regions to validate the model. Here we present the recall values for every CTCF, BEAF-32 and su(Hw) data set used in this manuscript; see Table S1 in *Appendix*. For each individual plot, we considered the cases of No accessibility, Continuous accessibility and DHS accessibility.

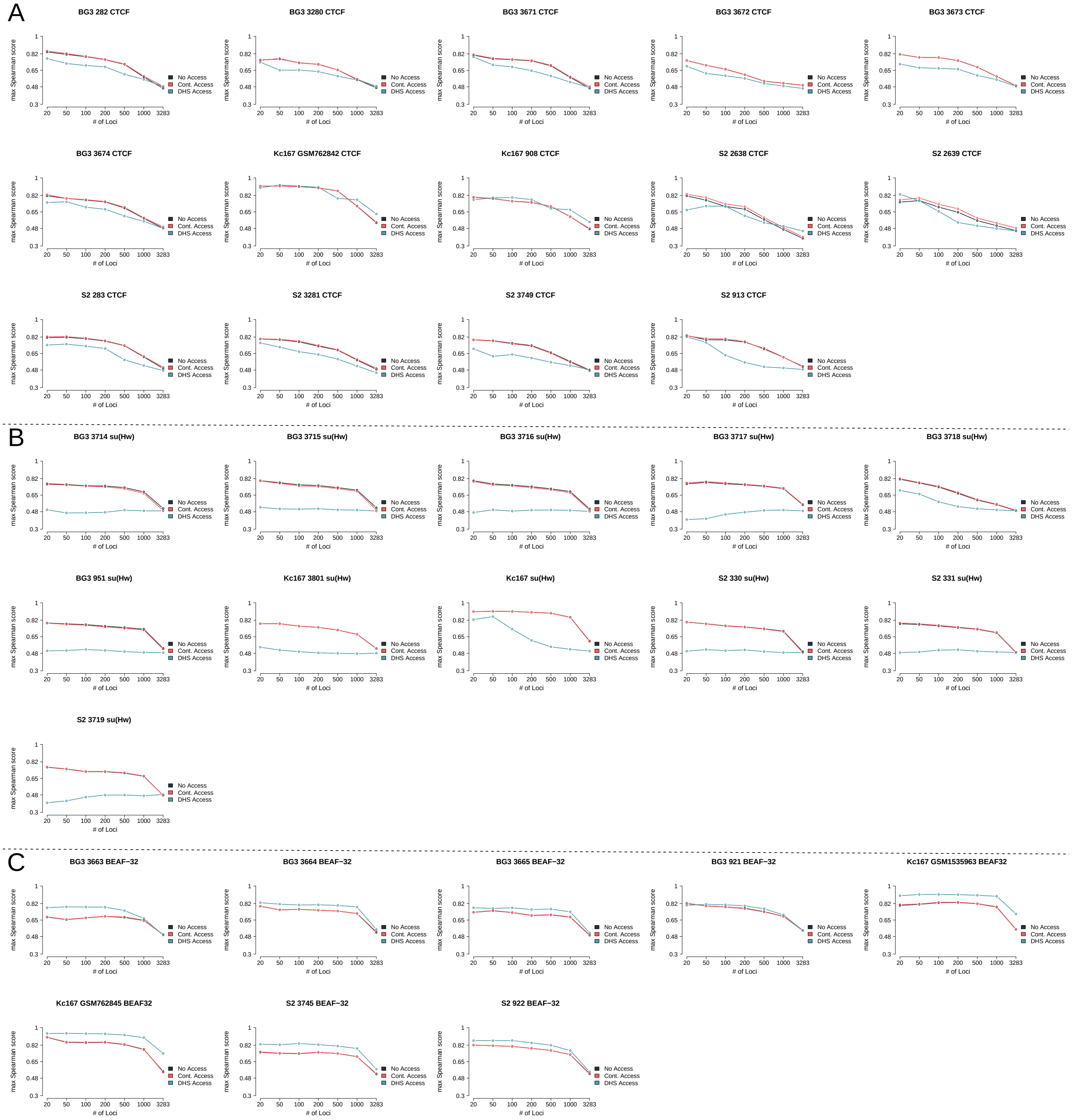


Figure S10: **Changes in spearman correlation as the number of regions used for validation increases** We selected varying numbers of regions to validate the model. Here we present the spearman correlation values for every CTCF, BEAF-32 and su(Hw) data set used in this manuscript; see Table S1 in *Appendix*. For each individual plot, we considered the cases of No accessibility,Continuous accessibility and DHS accessibility.

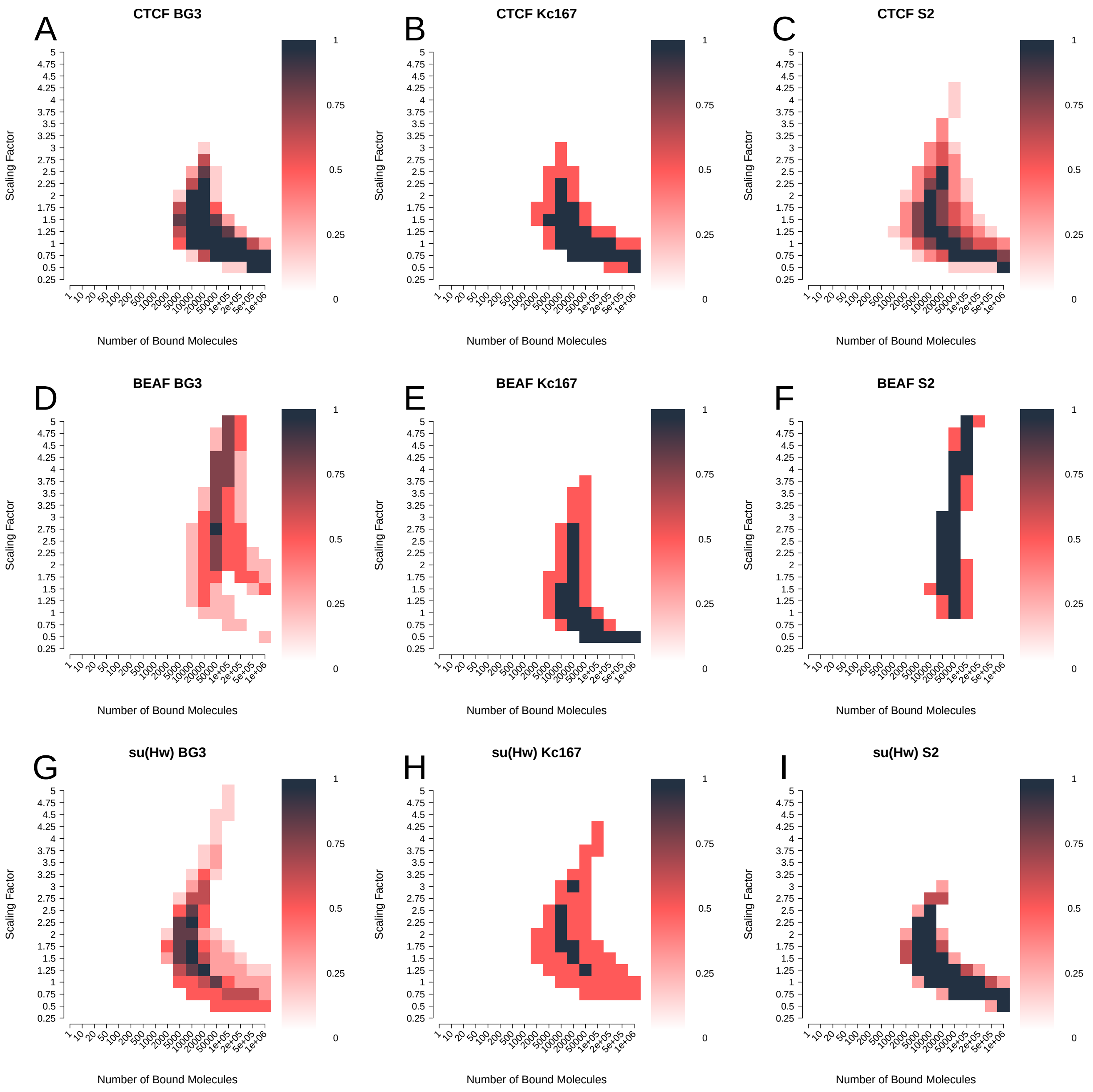


Figure S11: **Optimal parameters consistency among biological replicates for MSE using DHS accessibility.** Heat maps show an overlay of the top 10 % combinations of parameters when minimising MSE for: (**A-C**) CTCF, (**D-F**) BEAF-32 and (**G-I**) su(Hw). We plot the following cell lines: (**A**, **D** and **G**) BG3, (**B**, **E** and **H**) Kc167 and (**C**, **F** and **I**) BG3.

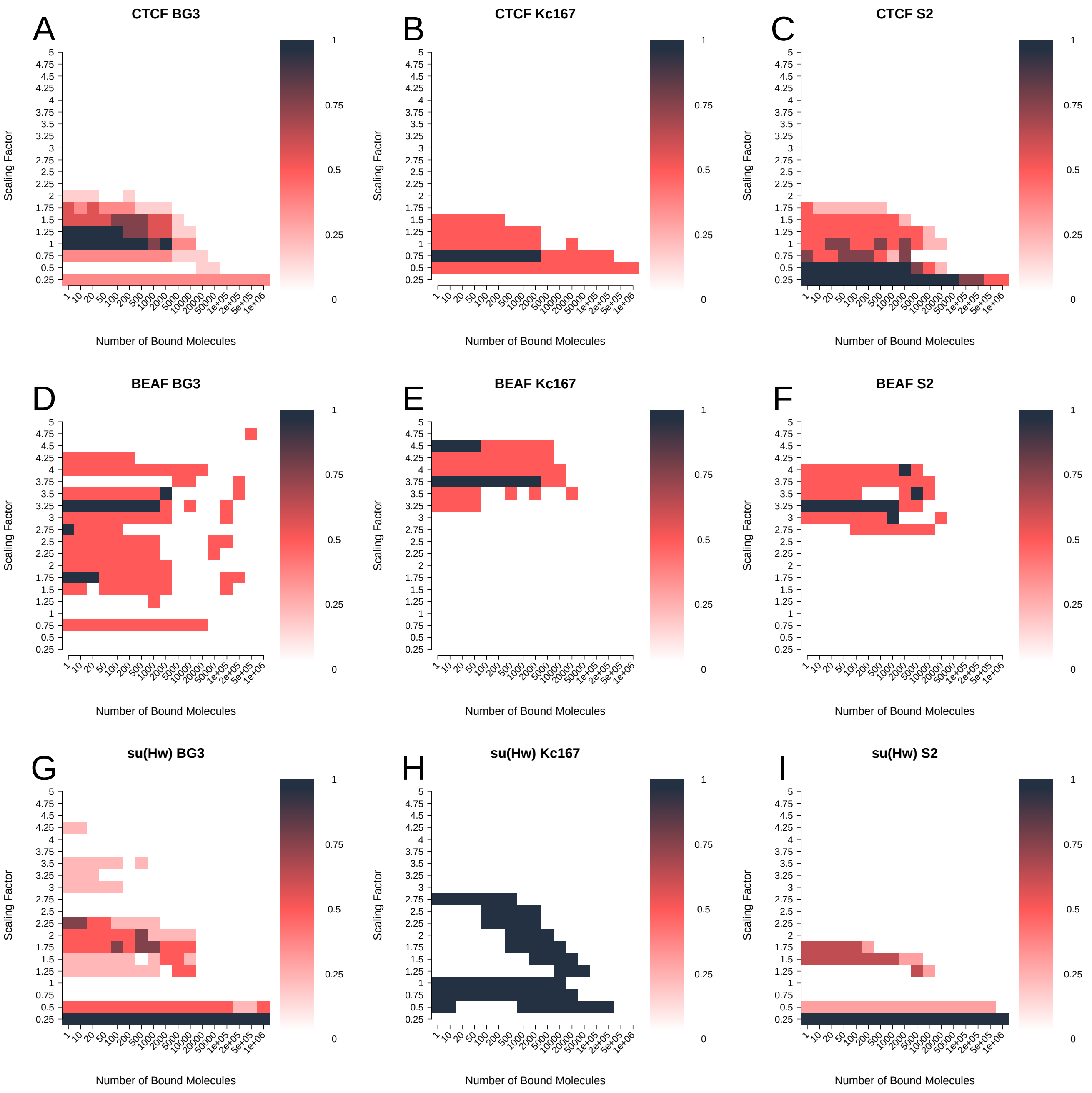


Figure S12: **Optimal parameters consistency among biological replicates for AUC using DHS accessibility.** Heat maps show an overlay of the top 10 % combinations of parameters when maximising AUC for: (**A-C**) CTCF, (**D-F**) BEAF-32 and (**G-I**) su(Hw). We plot the following cell lines: (**A**, **D** and **G**) BG3, (**B**, **E** and **H**) Kc167 and (**C**, **F** and **I**) BG3.

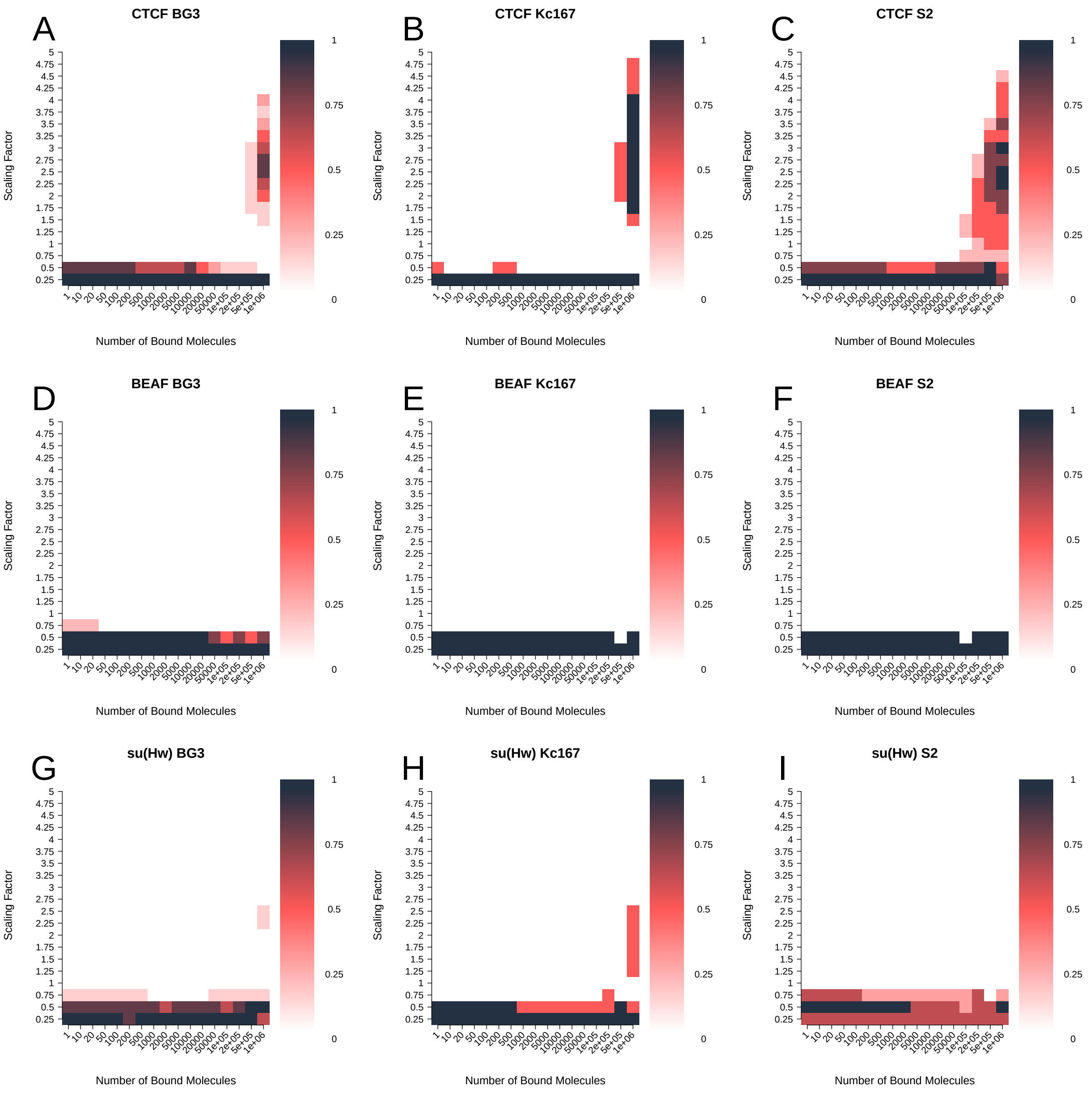


Figure S13: **Optimal parameters consistency among biological replicates for recall using DHS accessibility.** Heat maps show an overlay of the top 10 % combinations of parameters when maximising recall for: (**A-C**) CTCF, (**D-F**) BEAF-32 and (**G-I**) su(Hw). We plot the following cell lines: (**A**, **D** and **G**) BG3, (**B**, **E** and **H**) Kc167 and (**C**, **F** and **I**) BG3.

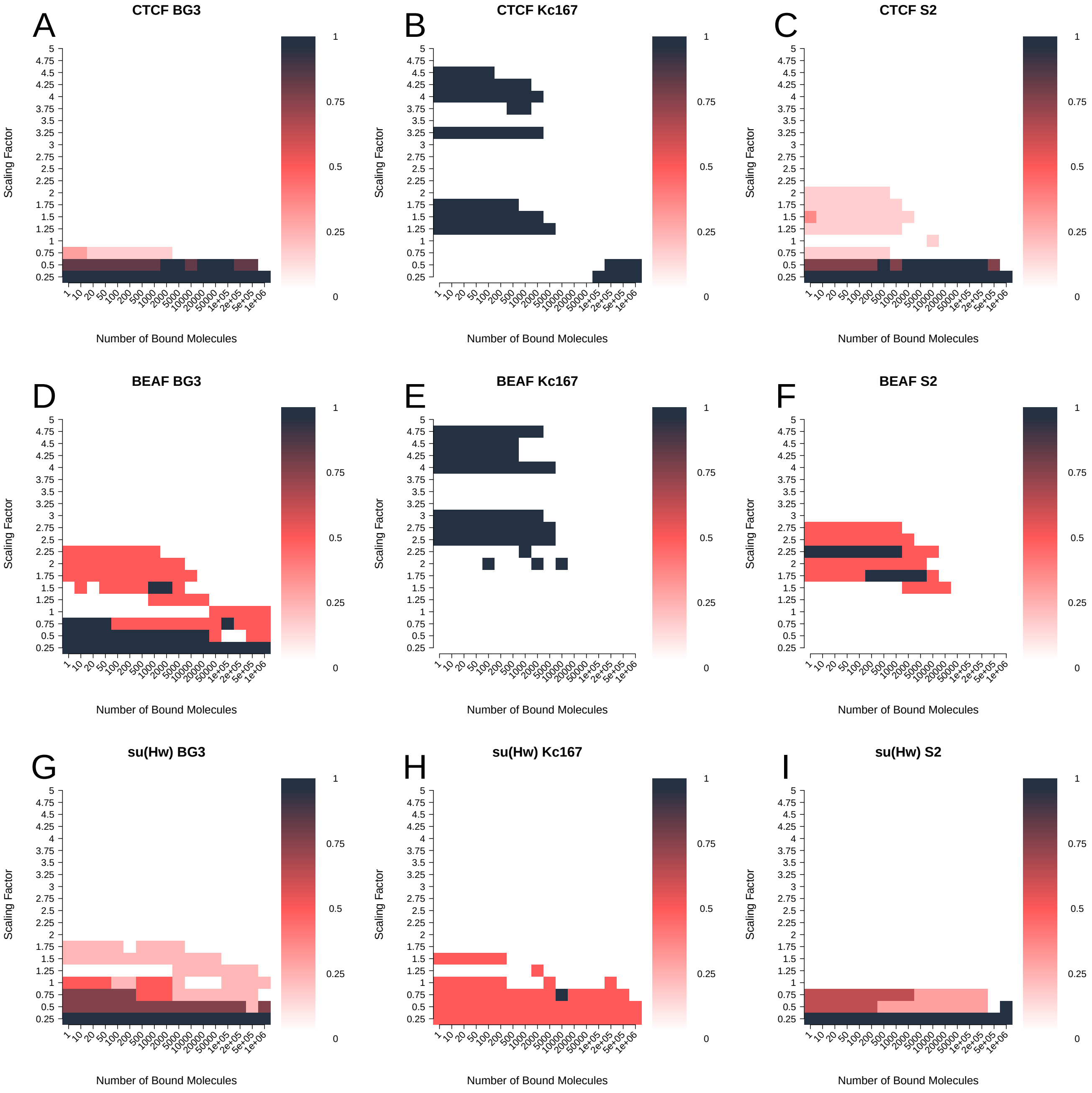


Figure S14: **Optimal parameters consistency among biological replicates for spearman correlation using DHS accessibility.** Heat maps show an overlay of the top 10 % combinations of parameters when maximising spearman correlation for: (**A-C**) CTCF, (**D-F**) BEAF-32 and (**G-I**) su(Hw). We plot the following cell lines: (**A**, **D** and **G**) BG3, (**B**, **E** and **H**) Kc167 and (**C**, **F** and **I**) BG3.
