## Supplemental Table for "Dissecting the binding mechanisms of transcription factors to DNA using a statistical thermodynamics framework"

**Table S1: Data sources for all TFs.** Data sets have been separated into modEncode ChIP data [1] , GEO ChIP data, Accessibility Data, and ENCODE data for *Homo sapiens* [1, 2]

| Cell Line | TF | modEncode Accession # |
|---|---|---|
| S2 | CTCF | 2638 |
| S2 | CTCF | 2639 |
| S2 | CTCF | 283 |
| S2 | CTCF | 3281 |
| S2 | CTCF | 3749 |
| S2 | CTCF | 913 |
| S2 | BEAF-32 | 922 |
| S2 | BEAF-32 | 3745 |
| S2 | BEAF-32 | 274 |
| S2 | Su(Hw) | 3719 |
| S2 | Su(Hw) | 330 |
| S2 | Su(Hw) | 331 |
| BG3 | CTCF | 282 |
| BG3 | CTCF | 3280 |
| BG3 | BEAF-32 | 3663 |
| BG3 | BEAF-32 | 3664 |
| BG3 | BEAF-32 | 3665 |
| BG3 | BEAF-32 | 921 |
| BG3 | CTCF | 3671 |
| BG3 | CTCF | 3672 |
| BG3 | CTCF | 3673 |
| BG3 | CTCF | 3674 |
| BG3 | Su(Hw) | 3714 |
| BG3 | Su(Hw) | 3715 |
| BG3 | Su(Hw) | 3716 |
| BG3 | Su(Hw) | 3717 |
| BG3 | Su(Hw) | 3718 |
| BG3 | Su(Hw) | 951 |
| Kc167 | CTCF | 908 |
| Kc167 | Su(Hw) | 3801 |

| Cell Line | TF | GEO Accession # |
|---|---|---|
| Kc167 | CTCF | GSM762842 |
| Kc167 | Su(Hw) | GSM762839 |
| Kc167 | BEAF-32 | GSM762845 |
| Kc167 | BEAF-32 | GSM1535963 |

| Cell Line | Method | GEO Accession # |
|---|---|---|
| Kc167 | DNase | Kharchenko et al. 2011 |
| S2 | DNase | Kharchenko et al. 2011 |
| BG3 | DNase | Kharchenko et al. 2011 |
| Kc167 | ATAC-seq | GSE122575 |

| Cell Line | Method | File Type | ENCODE # |
|---|---|---|---|
| Astrocyte | DNase | BAM | ENCFF384CCQ |
| Astrocyte | DNase | BAM | ENCFF885IAD |
| Astrocyte | DNase | BigWig | ENCFF901UBX |
| Astrocyte | DNase | bed | ENCFF021SAS |
| Astrocyte | DNase | bed | ENCFF352LYZ |
| Astrocyte | ChIP-seq | bigWig | ENCFF424JNY |
| Astrocyte | ChIP-seq | bed | ENCFF600CYD |
| Astrocyte | ChIP-seq | bed | ENCFF183YLB |

**Table S2: Goodness of Fit metrics:** ChIPanalyser offers 12 goodness of fit metrics grouped into two classes *Dissimilarity* and *Similarity.* Symbolically each metric is either a measure of how different two datasets are (Dissimilarity) or a measure of how similar two datasets are (Similarity). TP - true positives, TN - true negatives, FP - false positives and FN - false negatives. MCC represents Matthews Correlation Coefficient.

| Metric | Description | Type |
|---|---|---|
| MSE | Mean Squared Error | Dissimilarity |
| K-S distance | Kolmogorov-Smirnov Goodness-of-Fit Test | Dissimilarity |
| Geometric ratio | $\frac{\int_a^b \lvert f(x)-g(x) \rvert dx}{\int_a^b min(f(x),g(x))dx}$ | Dissimilarity |
| Recall | $\frac{TP}{TP+FN}$ | Dissimilarity |
| Pearson | Pearson correlation coefficient between predicted and ChIP profiles | Similarity |
| Spearman | Spearman correlation coefficient between predicted and ChIP profiles | Similarity |
| Kendall | Kendall correlation coefficient between predicted and ChIP profiles | Similarity |
| Precision | $\frac{TP}{TP+FP}$ | Similarity |
| F-score | $\frac{2TP}{2TP+FP+FN}$ | Similarity |
| Accuracy | $\frac{TP+TN}{TP+TN+FP+FN}$ | Similarity |
| MCC | $\frac{TP \times TN - FP \times FN}{\sqrt{(TP+FP)(TP+FN)(TN+FP)(TN+FN)}}$ | Similarity |
| AUC | Area Under the ROC (Receiver Operator Characteristic ) Curve | Similarity |

**Table S3: Optimal set of parameters after training on top ten regions and minimising MSE** From left to right, the white column shows the optimal parameters selected and associated MSE during training when No Accessibility was considered. Light grey represents optimal parameters selected and associated MSE during training when Continuous DNA accessibility scores were considered. Finally, dark grey are the optimal parameters selected and associated MSE when only DNase hypersensitivity sites (DHS) were considered.

| TF | # Bound | Lambda | MSE | # Bound | Lambda | MSE | # Bound | Lambda | MSE |
|---|---|---|---|---|---|---|---|---|---|
| BG3 modEncode 282 CTCF | 1e+06 | 0.75 | 0.009 | 1e+06 | 0.5 | 0.009 | 5e+05 | 0.75 | 0.009 |
| BG3 modEncode 3280 CTCF | 5e+05 | 0.75 | 0.013 | 5e+05 | 0.5 | 0.013 | 20000 | 1.5 | 0.018 |
| BG3 modEncode 3671 CTCF | 1e+06 | 0.75 | 0.008 | 1e+06 | 0.5 | 0.008 | 5e+05 | 0.75 | 0.007 |
| BG3 modEncode 3672 CTCF | 1e+06 | 0.5 | 0.006 | 50000 | 0.5 | 0.006 | 1e+05 | 1 | 0.007 |
| BG3 modEncode 3673 CTCF | 5e+05 | 0.75 | 0.01 | 20000 | 0.75 | 0.01 | 5e+05 | 0.75 | 0.008 |
| BG3 modEncode 3674 CTCF | 1e+06 | 0.75 | 0.01 | 1e+06 | 0.5 | 0.01 | 5e+05 | 0.75 | 0.011 |
| Kc167 GSM762842 CTCF | 50000 | 1 | 0.007 | 500 | 1 | 0.007 | 10000 | 1.5 | 0.006 |
| Kc167 modEncode 908 CTCF | 5e+05 | 0.75 | 0.011 | 50000 | 0.5 | 0.01 | 5e+05 | 0.75 | 0.009 |
| S2 modEncode 2638 CTCF | 50000 | 1.25 | 0.002 | 500 | 1.25 | 0.002 | 5000 | 1.5 | 0.002 |
| S2 modEncode 2639 CTCF | 20000 | 1.25 | 0.002 | 200 | 1.25 | 0.002 | 5000 | 2 | 0.002 |
| S2 modENCODE 283 CTCF | 5e+05 | 0.75 | 0.014 | 20000 | 0.5 | 0.013 | 1e+06 | 0.75 | 0.013 |
| S2 modEncode 3281 CTCF | 5e+05 | 0.75 | 0.017 | 20000 | 0.5 | 0.017 | 50000 | 2 | 0.019 |
| S2 modEncode 3749 CTCF | 1e+05 | 1 | 0.009 | 1000 | 1 | 0.009 | 20000 | 1.25 | 0.01 |
| S2 modEncode 913 CTCF | 2e+05 | 0.75 | 0.01 | 1e+06 | 0.25 | 0.01 | 5e+05 | 0.5 | 0.014 |
| BG3 modEncode 3714 Su(Hw) | 1e+05 | 2 | 0.023 | 5000 | 2 | 0.023 | 5000 | 2 | 0.045 |
| BG3 modEncode 3715 Su(Hw) | 1e+05 | 2 | 0.024 | 5000 | 2 | 0.024 | 10000 | 2.75 | 0.046 |
| BG3 modEncode 3716 Su(Hw) | 2e+05 | 1.5 | 0.016 | 5000 | 1.25 | 0.016 | 10000 | 2 | 0.031 |
| BG3 modEncode 3717 Su(Hw) | 1e+05 | 2 | 0.016 | 2000 | 1.75 | 0.016 | 10000 | 2.25 | 0.032 |
| BG3 modEncode 3718 Su(Hw) | 50000 | 1.75 | 0.012 | 1000 | 1.75 | 0.012 | 5e+05 | 1 | 0.019 |
| BG3 modEncode 951 Su(Hw) | 1e+05 | 1.75 | 0.017 | 2000 | 1.5 | 0.017 | 2e+05 | 0.75 | 0.041 |
| Kc167 modEncode 3801 Su(Hw) | 1e+05 | 2.25 | 0.017 | 5000 | 2.25 | 0.017 | 2e+05 | 1.25 | 0.03 |
| Kc167 Su(Hw) | 50000 | 1.25 | 0.004 | 200 | 1.25 | 0.004 | 10000 | 1.25 | 0.005 |
| S2 modEncode 330 Su(Hw) | 2e+05 | 1.75 | 0.022 | 5000 | 1.75 | 0.022 | 5e+05 | 0.75 | 0.05 |
| S2 modEncode 331 Su(Hw) | 1e+06 | 1.25 | 0.017 | 5000 | 1.25 | 0.017 | 5e+05 | 0.75 | 0.037 |
| S2 modEncode 3719 Su(Hw) | 2e+05 | 2.25 | 0.033 | 10000 | 2.25 | 0.032 | 2e+05 | 0.75 | 0.066 |
| BG3 modEncode 3663 BEAF-32 | 1e+05 | 2.75 | 0.026 | 10000 | 2.5 | 0.026 | 50000 | 3 | 0.023 |
| BG3 modEncode 3664 BEAF-32 | 1e+05 | 2.5 | 0.027 | 10000 | 2.5 | 0.027 | 1e+05 | 4 | 0.019 |
| BG3 modEncode 3665 BEAF-32 | 2e+05 | 3 | 0.039 | 20000 | 2.75 | 0.04 | 2e+05 | 2.5 | 0.034 |
| BG3 modEncode 921 BEAF-32 | 20000 | 1.25 | 0.012 | 2000 | 1.75 | 0.012 | 20000 | 2.5 | 0.008 |
| Kc167 GSM1535963 BEAF32 | 50000 | 0.75 | 0.012 | 1000 | 1 | 0.012 | 50000 | 0.75 | 0.008 |
| Kc167 GSM762845 BEAF32 | 20000 | 1 | 0.006 | 1000 | 1.25 | 0.006 | 10000 | 1.25 | 0.005 |
| S2 modEncode 3745 BEAF-32 | 50000 | 1.25 | 0.017 | 1000 | 1 | 0.017 | 50000 | 3.25 | 0.014 |
| S2 modEncode 922 BEAF-32 | 50000 | 1.5 | 0.014 | 2000 | 1.5 | 0.014 | 50000 | 3 | 0.009 |

**Table S4: Optimal set of parameters after training on top ten regions and maximising AUC** From left to right, the white column shows the optimal parameters selected and associated AUC during training when No Accessibility was considered. Light grey represents optimal parameters selected and associated AUC during training when Continuous DNA accessibility scores were considered. Finally, dark grey are the optimal parameters selected and associated AUC when only DNase hypersensitivity sites (DHS) were considered.

| TF | # Bound | Lambda | AUC | # Bound | Lambda | AUC | # Bound | Lambda | AUC |
|---|---|---|---|---|---|---|---|---|---|
| BG3 modEncode 282 CTCF | 5e+05 | 0.25 | 0.944 | 20000 | 0.25 | 0.943 | 1 | 0.25 | 0.885 |
| BG3 modEncode 3280 CTCF | 20000 | 0.25 | 0.895 | 2000 | 0.25 | 0.895 | 5e+05 | 0.25 | 0.812 |
| BG3 modEncode 3671 CTCF | 2e+05 | 0.5 | 0.936 | 5e+05 | 0.25 | 0.937 | 10000 | 1 | 0.946 |
| BG3 modEncode 3672 CTCF | 10 | 1 | 0.922 | 1 | 1 | 0.922 | 2000 | 1.25 | 0.832 |
| BG3 modEncode 3673 CTCF | 20000 | 0.5 | 0.903 | 1000 | 0.5 | 0.903 | 20 | 1.25 | 0.853 |
| BG3 modEncode 3674 CTCF | 1e+06 | 0.25 | 0.929 | 5e+05 | 0.25 | 0.929 | 500 | 0.75 | 0.847 |
| Kc167 GSM762842 CTCF | 1e+06 | 0.5 | 0.954 | 10000 | 0.5 | 0.955 | 1e+06 | 0.5 | 0.887 |
| Kc167 modEncode 908 CTCF | 1e+05 | 0.25 | 0.952 | 50 | 0.25 | 0.952 | 1 | 1.25 | 0.963 |
| S2 modEncode 2638 CTCF | 50000 | 0.5 | 0.99 | 100 | 0.5 | 0.99 | 5000 | 0.5 | 0.963 |
| S2 modEncode 2639 CTCF | 100 | 2 | 0.915 | 1 | 2 | 0.916 | 2e+05 | 0.25 | 0.857 |
| S2 modENCODE 283 CTCF | 1e+06 | 0.25 | 0.926 | 200 | 0.25 | 0.926 | 10 | 1.25 | 0.886 |
| S2 modEncode 3281 CTCF | 1e+06 | 0.25 | 0.882 | 10000 | 0.25 | 0.882 | 50 | 1 | 0.864 |
| S2 modEncode 3749 CTCF | 50000 | 0.25 | 0.857 | 10 | 0.25 | 0.857 | 1e+06 | 0.25 | 0.848 |
| S2 modEncode 913 CTCF | 2000 | 0.75 | 0.938 | 1 | 0.75 | 0.938 | 2000 | 0.5 | 0.872 |
| BG3 modEncode 3714 Su(Hw) | 20000 | 0.75 | 0.928 | 100 | 0.75 | 0.928 | 1 | 0.25 | 0.458 |
| BG3 modEncode 3715 Su(Hw) | 50 | 0.75 | 0.927 | 1 | 0.75 | 0.927 | 1 | 0.25 | 0.606 |
| BG3 modEncode 3716 Su(Hw) | 10000 | 0.75 | 0.929 | 50 | 0.75 | 0.93 | 1 | 0.25 | 0.677 |
| BG3 modEncode 3717 Su(Hw) | 5000 | 0.75 | 0.927 | 20 | 0.75 | 0.927 | 10000 | 1.25 | 0.662 |
| BG3 modEncode 3718 Su(Hw) | 1 | 0.75 | 0.93 | 10 | 0.75 | 0.93 | 20 | 1.5 | 0.766 |
| BG3 modEncode 951 Su(Hw) | 2000 | 0.75 | 0.931 | 10 | 0.75 | 0.931 | 200 | 0.25 | 0.583 |
| Kc167 modEncode 3801 Su(Hw) | 50000 | 0.75 | 0.925 | 20 | 0.75 | 0.925 | 20000 | 0.75 | 0.548 |
| Kc167 Su(Hw) | 1e+06 | 0.75 | 0.962 | 500 | 0.75 | 0.962 | 20000 | 1.25 | 0.915 |
| S2 modEncode 330 Su(Hw) | 50000 | 0.5 | 0.931 | 50 | 0.5 | 0.931 | 1 | 0.25 | 0.565 |
| S2 modEncode 331 Su(Hw) | 1e+05 | 0.5 | 0.937 | 100 | 0.75 | 0.937 | 5000 | 0.25 | 0.643 |
| S2 modEncode 3719 Su(Hw) | 10000 | 1 | 0.917 | 50 | 1 | 0.916 | 1 | 0.25 | 0.43 |
| BG3 modEncode 3663 BEAF-32 | 20000 | 0.25 | 0.66 | 1000 | 0.25 | 0.661 | 1 | 2.25 | 0.81 |
| BG3 modEncode 3664 BEAF-32 | 5000 | 4 | 0.789 | 1000 | 4 | 0.789 | 100 | 3.5 | 0.859 |
| BG3 modEncode 3665 BEAF-32 | 500 | 1.5 | 0.726 | 1 | 1.5 | 0.726 | 10000 | 0.75 | 0.816 |
| BG3 modEncode 921 BEAF-32 | 5000 | 4.25 | 0.831 | 1000 | 4 | 0.831 | 1e+05 | 3 | 0.897 |
| Kc167 GSM1535963 BEAF32 | 2e+05 | 0.25 | 0.847 | 5000 | 0.25 | 0.847 | 1000 | 3.75 | 0.963 |
| Kc167 GSM762845 BEAF32 | 10000 | 4 | 0.937 | 1000 | 4 | 0.937 | 50 | 4.25 | 0.986 |
| S2 modEncode 3745 BEAF-32 | 2e+05 | 0.25 | 0.857 | 1000 | 0.25 | 0.857 | 200 | 3.75 | 0.869 |
| S2 modEncode 922 BEAF-32 | 10000 | 4 | 0.825 | 2000 | 4.25 | 0.826 | 1 | 3 | 0.928 |

**Table S5: Optimal set of parameters after training on top ten regions and maximising recall** From left to right, the white column shows the optimal parameters selected and associated recall during training when No Accessibility was considered. Light grey represents optimal parameters selected and associated recall during training when Continuous DNA accessibility scores were considered. Finally, dark grey are the optimal parameters selected and associated recall when only DNase hypersensitivity sites (DHS) were considered.

| TF | # Bound | Lambda | Recall | # Bound | Lambda | Recall | # Bound | Lambda | Recall |
|---|---|---|---|---|---|---|---|---|---|
| BG3 modEncode 282 CTCF | 20000 | 0.75 | 0.887 | 1000 | 0.75 | 0.887 | 1 | 1.5 | 0.778 |
| BG3 modEncode 3280 CTCF | 10000 | 0.75 | 0.816 | 500 | 0.75 | 0.818 | 1 | 1.25 | 0.676 |
| BG3 modEncode 3671 CTCF | 10000 | 0.75 | 0.884 | 500 | 0.75 | 0.884 | 1000 | 1.5 | 0.859 |
| BG3 modEncode 3672 CTCF | 1 | 1.25 | 0.883 | 1 | 1.25 | 0.883 | 5000 | 1.5 | 0.693 |
| BG3 modEncode 3673 CTCF | 20000 | 0.75 | 0.855 | 500 | 0.75 | 0.854 | 100 | 1.5 | 0.758 |
| BG3 modEncode 3674 CTCF | 10000 | 0.75 | 0.865 | 2000 | 0.75 | 0.865 | 100 | 1.5 | 0.735 |
| Kc167 GSM762842 CTCF | 5e+05 | 0.75 | 0.932 | 5000 | 0.75 | 0.932 | 10000 | 1.75 | 0.807 |
| Kc167 modEncode 908 CTCF | 10000 | 0.75 | 0.88 | 50 | 0.75 | 0.88 | 1 | 1.25 | 0.831 |
| S2 modEncode 2638 CTCF | 10000 | 1 | 0.962 | 100 | 1 | 0.962 | 2000 | 1.5 | 0.859 |
| S2 modEncode 2639 CTCF | 100 | 2 | 0.891 | 1 | 2.25 | 0.892 | 2000 | 1.75 | 0.727 |
| S2 modENCODE 283 CTCF | 10000 | 0.75 | 0.86 | 50 | 0.75 | 0.86 | 1 | 1.5 | 0.744 |
| S2 modEncode 3281 CTCF | 20000 | 0.75 | 0.805 | 50 | 0.75 | 0.805 | 500 | 1.25 | 0.71 |
| S2 modEncode 3749 CTCF | 20000 | 1 | 0.816 | 20 | 1 | 0.814 | 2000 | 1.25 | 0.682 |
| S2 modEncode 913 CTCF | 2000 | 1 | 0.875 | 20 | 1 | 0.875 | 1 | 1.75 | 0.711 |
| BG3 modEncode 3714 Su(Hw) | 500 | 1.25 | 0.85 | 20 | 1.25 | 0.85 | 10000 | 1.25 | 0.208 |
| BG3 modEncode 3715 Su(Hw) | 1000 | 1.25 | 0.848 | 10 | 1.25 | 0.848 | 10 | 2.25 | 0.397 |
| BG3 modEncode 3716 Su(Hw) | 20000 | 1.25 | 0.86 | 50 | 1.25 | 0.86 | 10000 | 1.5 | 0.443 |
| BG3 modEncode 3717 Su(Hw) | 2000 | 1.25 | 0.855 | 20 | 1.25 | 0.855 | 10000 | 1.25 | 0.419 |
| BG3 modEncode 3718 Su(Hw) | 5000 | 1.25 | 0.869 | 50 | 1.25 | 0.868 | 1 | 2 | 0.547 |
| BG3 modEncode 951 Su(Hw) | 5000 | 1.25 | 0.846 | 50 | 1.25 | 0.846 | 1 | 4.25 | 0.346 |
| Kc167 modEncode 3801 Su(Hw) | 10 | 1.25 | 0.851 | 1 | 1.25 | 0.851 | 20 | 1.5 | 0.225 |
| Kc167 Su(Hw) | 50000 | 1 | 0.936 | 50 | 1 | 0.935 | 20000 | 1.25 | 0.833 |
| S2 modEncode 330 Su(Hw) | 5000 | 1.25 | 0.841 | 20 | 1.25 | 0.841 | 5000 | 1.5 | 0.286 |
| S2 modEncode 331 Su(Hw) | 5000 | 1.25 | 0.863 | 20 | 1.25 | 0.862 | 10000 | 1.25 | 0.406 |
| S2 modEncode 3719 Su(Hw) | 5000 | 1.25 | 0.813 | 10 | 1.25 | 0.812 | 1e+05 | 1 | 0.132 |
| BG3 modEncode 3663 BEAF-32 | 10 | 1 | 0.627 | 20 | 1 | 0.627 | 1 | 2.75 | 0.69 |
| BG3 modEncode 3664 BEAF-32 | 5000 | 4 | 0.762 | 1000 | 4 | 0.761 | 5000 | 4 | 0.732 |
| BG3 modEncode 3665 BEAF-32 | 2000 | 2 | 0.687 | 200 | 2 | 0.687 | 2e+05 | 1.75 | 0.663 |
| BG3 modEncode 921 BEAF-32 | 5000 | 4.25 | 0.807 | 1000 | 4.25 | 0.806 | 1e+05 | 3 | 0.829 |
| Kc167 GSM1535963 BEAF32 | 1 | 1.5 | 0.823 | 1 | 1.5 | 0.823 | 1000 | 3.75 | 0.902 |
| Kc167 GSM762845 BEAF32 | 10000 | 4 | 0.918 | 1000 | 4 | 0.918 | 50 | 4.25 | 0.95 |
| S2 modEncode 3745 BEAF-32 | 1000 | 0.75 | 0.806 | 20 | 0.75 | 0.806 | 1 | 3.25 | 0.745 |
| S2 modEncode 922 BEAF-32 | 10000 | 4 | 0.799 | 2000 | 4.25 | 0.801 | 2000 | 3.25 | 0.856 |

**Table S6: Optimal set of parameters after training on top ten regions and maximising Spearman correlation** From left to right, the white column shows the optimal parameters selected and associated spearman correlation during training when No Accessibility was considered. Light grey represents optimal parameters selected and associated Spearman correlation during training when Continuous DNA accessibility scores were considered. Finally, dark grey are the optimal parameters selected and associated Spearman correlation when only DNase hypersensitivity sites (DHS) were considered.

| TF | # Bound | Lambda | spearman | # Bound | Lambda | spearman | # Bound | Lambda | spearman |
|---|---|---|---|---|---|---|---|---|---|
| BG3 modEncode 282 CTCF | 1e+06 | 0.25 | 0.584 | 2e+05 | 0.25 | 0.584 | 5e+05 | 0.25 | 0.523 |
| BG3 modEncode 3280 CTCF | 1e+06 | 0.25 | 0.512 | 2e+05 | 0.25 | 0.513 | 5e+05 | 0.25 | 0.461 |
| BG3 modEncode 3671 CTCF | 1e+06 | 0.25 | 0.582 | 1e+06 | 0.25 | 0.583 | 1 | 0.25 | 0.581 |
| BG3 modEncode 3672 CTCF | 1e+05 | 0.5 | 0.502 | 5000 | 0.5 | 0.503 | 1 | 0.25 | 0.404 |
| BG3 modEncode 3673 CTCF | 1e+06 | 0.25 | 0.463 | 1e+06 | 0.25 | 0.464 | 1 | 0.25 | 0.462 |
| BG3 modEncode 3674 CTCF | 1e+06 | 0.25 | 0.483 | 50000 | 0.25 | 0.483 | 2000 | 0.5 | 0.471 |
| Kc167 GSM762842 CTCF | 1 | 0.25 | 0.299 | 5e+05 | 0.25 | 0.312 | 20 | 3.25 | 0.367 |
| Kc167 modEncode 908 CTCF | 5e+05 | 0.25 | 0.551 | 100 | 0.25 | 0.551 | 10 | 1.25 | 0.638 |
| S2 modEncode 2638 CTCF | 1e+06 | 0.25 | 0.608 | 10000 | 0.25 | 0.608 | 5e+05 | 0.25 | 0.544 |
| S2 modEncode 2639 CTCF | 1 | 0.25 | 0.438 | 10000 | 0.25 | 0.438 | 1 | 0.25 | 0.323 |
| S2 modENCODE 283 CTCF | 1 | 0.25 | 0.596 | 20000 | 0.25 | 0.597 | 1 | 0.25 | 0.517 |
| S2 modEncode 3281 CTCF | 1e+06 | 0.25 | 0.496 | 500 | 0.25 | 0.496 | 5e+05 | 0.25 | 0.448 |
| S2 modEncode 3749 CTCF | 1e+06 | 0.25 | 0.446 | 10000 | 0.25 | 0.448 | 5e+05 | 0.25 | 0.467 |
| S2 modEncode 913 CTCF | 5e+05 | 0.25 | 0.613 | 10000 | 0.25 | 0.614 | 200 | 1.25 | 0.556 |
| BG3 modEncode 3714 Su(Hw) | 5e+05 | 0.5 | 0.689 | 1000 | 0.5 | 0.689 | 1 | 0.25 | -0.109 |
| BG3 modEncode 3715 Su(Hw) | 50 | 0.75 | 0.704 | 1 | 0.75 | 0.704 | 1 | 0.25 | 0.1 |
| BG3 modEncode 3716 Su(Hw) | 1000 | 0.75 | 0.705 | 1 | 0.75 | 0.705 | 1 | 0.25 | 0.22 |
| BG3 modEncode 3717 Su(Hw) | 2000 | 0.75 | 0.682 | 1 | 0.75 | 0.682 | 5e+05 | 1 | 0.238 |
| BG3 modEncode 3718 Su(Hw) | 1 | 0.5 | 0.647 | 50 | 0.5 | 0.645 | 1 | 0.75 | 0.462 |
| BG3 modEncode 951 Su(Hw) | 5e+05 | 0.5 | 0.718 | 20000 | 0.5 | 0.721 | 1 | 0.25 | 0.056 |
| Kc167 modEncode 3801 Su(Hw) | 1e+06 | 0.5 | 0.674 | 200 | 0.5 | 0.674 | 1e+05 | 0.75 | 0.062 |
| Kc167 Su(Hw) | 50000 | 0.75 | 0.421 | 20 | 0.75 | 0.428 | 200 | 1.5 | 0.361 |
| S2 modEncode 330 Su(Hw) | 1e+06 | 0.5 | 0.748 | 1e+06 | 0.25 | 0.757 | 1 | 0.25 | 0.037 |
| S2 modEncode 331 Su(Hw) | 5e+05 | 0.5 | 0.675 | 1e+06 | 0.25 | 0.691 | 1 | 0.25 | 0.194 |
| S2 modEncode 3719 Su(Hw) | 50000 | 0.75 | 0.743 | 100 | 0.75 | 0.746 | 1 | 0.25 | -0.266 |
| BG3 modEncode 3663 BEAF-32 | 100 | 0.5 | 0.16 | 1 | 0.5 | 0.16 | 500 | 0.5 | 0.503 |
| BG3 modEncode 3664 BEAF-32 | 500 | 0.25 | 0.317 | 20 | 0.25 | 0.317 | 1000 | 1.75 | 0.507 |
| BG3 modEncode 3665 BEAF-32 | 50000 | 0.25 | 0.341 | 1 | 0.75 | 0.34 | 50000 | 0.75 | 0.512 |
| BG3 modEncode 921 BEAF-32 | 1e+06 | 0.25 | 0.252 | 1e+05 | 0.25 | 0.253 | 1000 | 0.5 | 0.31 |
| Kc167 GSM1535963 BEAF32 | 2e+05 | 0.25 | 0.385 | 2000 | 0.25 | 0.385 | 1000 | 3 | 0.575 |
| Kc167 GSM762845 BEAF32 | 1e+06 | 0.25 | 0.297 | 10000 | 0.25 | 0.296 | 200 | 4 | 0.552 |
| S2 modEncode 3745 BEAF-32 | 50000 | 0.25 | 0.385 | 200 | 0.25 | 0.385 | 50 | 2.25 | 0.517 |
| S2 modEncode 922 BEAF-32 | 5e+05 | 0.25 | 0.238 | 5000 | 0.25 | 0.237 | 5000 | 1.75 | 0.448 |
